# PLA2G15 is a Lysosomal BMP Hydrolase and its Targeting Ameliorates Lysosomal Disease

**DOI:** 10.1101/2024.06.07.597919

**Authors:** Kwamina Nyame, Jian Xiong, Hisham N. Alsohybe, Arthur PH de Jong, Isabelle V. Peña, Ricardo de Miguel, Thijn R. Brummelkamp, Guido Hartmann, Sebastian M.B. Nijman, Matthijs Raaben, Judith A. Simcox, Vincent A. Blomen, Monther Abu-Remaileh

## Abstract

Lysosomes catabolize lipids and other biological molecules, a function essential for cellular and organismal homeostasis. Key to lipid catabolism in the lysosome is bis(monoacylglycero)phosphate (BMP), a major lipid constituent of intralysosomal vesicles and a stimulator of lipid-degrading enzymes. BMP levels are altered in a broad spectrum of human conditions, including neurodegenerative diseases. While a lysosomal BMP synthase was recently discovered, the enzymes that mediate BMP turnover has remained elusive. Here we show that the lysosomal phospholipase PLA2G15 is a physiological BMP hydrolase. We further demonstrate that BMP’s resistance to hydrolysis in the lysosome is conferred by the combination of its unique *sn2, sn2’* esterification position and stereochemistry, as neither feature alone is sufficient to provide this resistance. Purified PLA2G15 catabolizes most BMP species derived from cell and tissue lysosomes under acidic conditions. Furthermore, PLA2G15 catalytic activity against synthesized BMP stereoisomers with primary esters was comparable to its canonical substrates challenging the long-held thought that BMP’s unique stereochemistry is sufficient to confer resistance to acid phospholipases. Conversely, BMP with secondary esters and *S,S* stereoconfiguration is intrinsically stable in vitro and requires acyl migration for hydrolysis in lysosomes. Consistent with our biochemical data, PLA2G15-deficient cells and tissues accumulate multiple BMP species, a phenotype reversible by supplementing wildtype PLA2G15 but not its catalytically dead mutant. In addition, targeting PLA2G15 to increase BMP reverses the cholesterol phenotype in Niemann Pick Disease Type C (NPC1) patient fibroblasts and significantly ameliorates disease pathologies in NPC1-deficient mice leading to extended lifespan. Our findings establish the rules that govern the stability of BMP in the lysosome and identify PLA2G15 as a lysosomal BMP hydrolase and a potential target for therapeutic intervention in neurodegenerative diseases.

## Main

Lysosomes are vital for cellular waste removal, nutrient recycling, and maintaining homeostasis, particularly through breaking down complex lipids using hydrolytic enzymes. Lipid degradation within lysosomes is facilitated by intralysosomal vesicles rich in a unique lipid called bis(monoacylglycero)phosphate (BMP) ^1^. BMP significantly enhances lysosomal lipid metabolism, and its imbalance is a hallmark of various lysosome-associated diseases, including age-related neurodegeneration, viral infection, cancer, and atherosclerotic cardiovascular disease ^1–4^.

While we recently identified the lysosomal BMP synthase ^2^, which was then validated as the sole lysosomal BMP synthase in cells by another group that also identified potential circulating machineries that might make BMP ^5^, many questions remain on the degradation of BMPs. BMP is a structural isomer of phosphatidylglycerol (PG) with symmetrical acyl chain positions on the two glycerol moieties and a unique *S,S* stereoconfiguration (Fig. 1a) ^4,6^. Partial deacylation of these phospholipids generates one fatty acid and lysophosphatidylglycerol (LPG) (Fig. 1b), while its complete deacylation releases two fatty acids and glycerophosphorylglycerol (GPG) (Fig. 1c) ^7–10^. BMP is stable in the lysosomal environment, however, the rules that govern BMP catabolism remain unclear. Some reports speculate that its unique stereochemistry may confer resistance to its degradation by lysosomal hydrolases, a necessity for its proposed function in stimulating lipid degradation in the lysosome ^11,12^, while others suggest that BMP is susceptible to enzyme-mediated hydrolysis ^7,9,13–16^. Because of its therapeutic potential, understanding BMP turnover and uncovering its physiological hydrolases in the lysosome is of great interest. Here, we show that the unique *sn2, sn2’* position of acyl chains of BMP, in addition to its *S,S* stereoconfiguration, protects BMPs from lysosomal hydrolysis by the abundant lysosomal phospholipase A2, PLA2G15, which we establish as a BMP hydrolase in addition to its role as a general acid phospholipase B. In a parallel genetic screening effort, we identified PLA2G15 as a modifier of cholesterol staining and showed that knockdown of PLA2G15 reduces the cholesterol phenotype in NPC1 patient fibroblasts. In addition, PLA2G15 inactivation in the NPC1 mouse increases lifespan and reduces expression of key disease biomarkers. Thus, targeting PLA2G15 boosts BMP levels and holds therapeutic potential in NPC disease and possibly other lysosomal storage diseases.

**Fig. 1:**
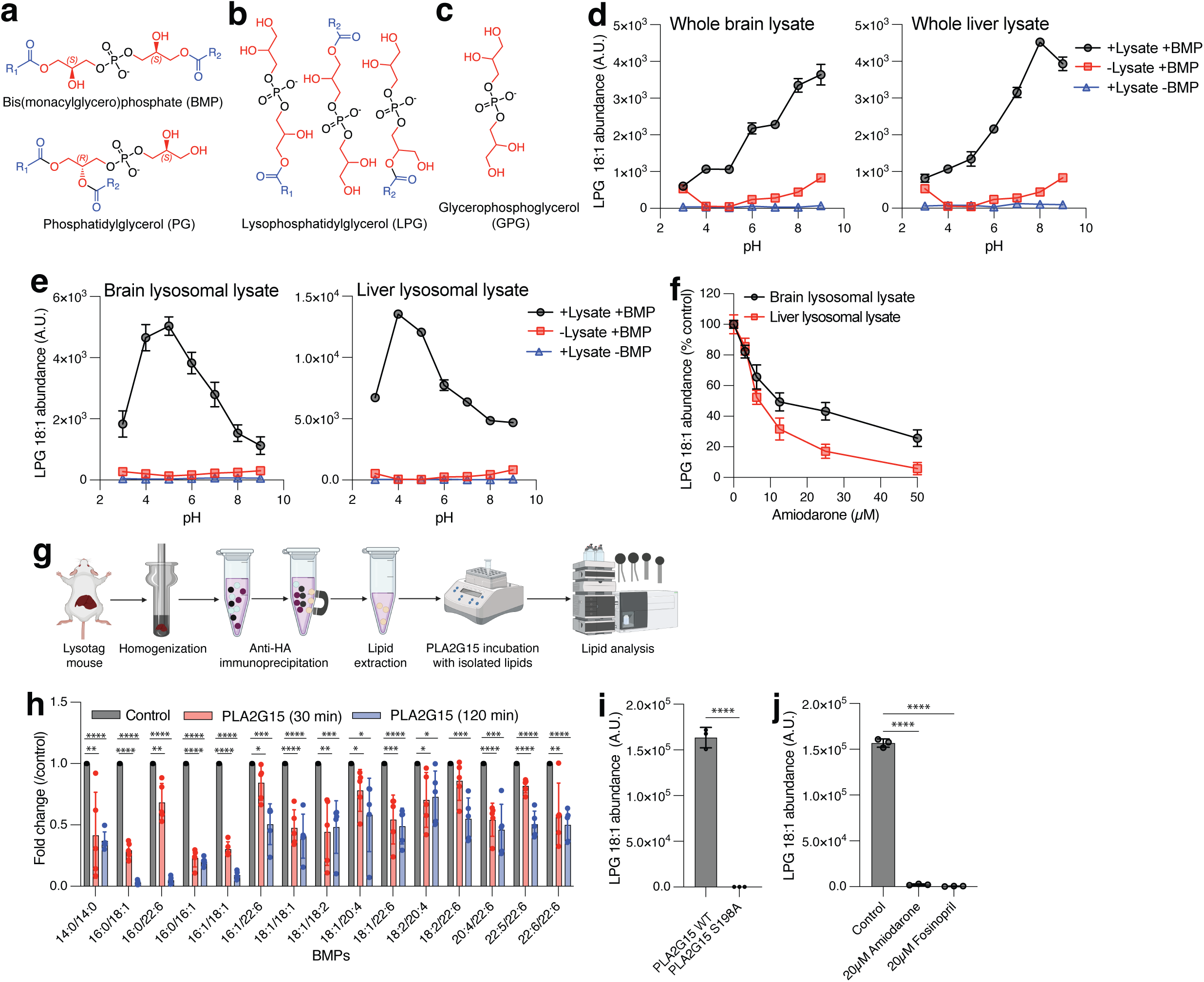
PLA2G15 hydrolyzes BMP lipids. **(a-c)** Bis(monoacylglycero)phosphate (BMP) is a structural isomer of phosphatidylglycerol (PG) with similar catabolic products. Chemical structures of BMP (top) and PG (bottom) in (a) and their degradation intermediates lysophosphatidylglycerol (LPG, all possible structures) in (b) and glycerophosphorylglycerol (GPG) in (c). Red represents two glycerol groups while blue represents the acyl chains. **(d)** BMP hydrolase activity in brain (left) and liver (right) lysates shows BMP is degraded across a broad pH spectrum including acidic environment. Hydrolysis of 3,3’ 18:1-*S,S* BMP (1 µM) in buffers with pH range between 3.0 and 9.0 (see methods). Controls had similar reaction buffers except with no lysate or BMP. Data are mean ± SD. *n* = 3 biological replicates. **(e)** Lysosomal lysates hydrolyze BMP with an acidic optimum. The same experiment as in (d) but using brain lysosomal lysate (left) and liver lysosomal lysate (right). **(f)** Amiodarone inhibits BMP lysosomal hydrolase activity in a concentration dependent manner. Activity assays were performed using 3,3’ 18:1-*S,S* BMP (1 µM) under acidic conditions (pH=5.0). Data are mean ± SD. *n* = 3 biological replicates. **(g and h)** PLA2G15 hydrolyzes BMPs isolated from mouse liver lysosomes. **(g)** Depiction of experimental design showing LysoIP to collect lysosomal lipids and incubation with PLA2G15 to monitor phospholipid abundances for 30 min or 120 min reactions. **(h)** Fold changes in the abundance of measured BMPs and each time point is compared to its control. **(i)** PLA2G15 S198A mutation completely abolishes BMP hydrolase activity. Hydrolysis of 1 µM of 3,3’ 18:1-*S,S* BMP by 100 nM recombinant PLA2G15 wildtype and S198A mutant under acidic conditions for 30 sec reaction time. BMP hydrolase activity was determined using LC-MS by calculating the abundance of LPG intermediate. Data are mean ± SD of *n* = 3 independent replicates from each protein. *p < 0.05, **p < 0.01, ***p < 0.001, ****p < 0.0001. **(j)** PLA2G15 inhibitors attenuate BMP hydrolase activity. The same experiment as in (i) using wildtype PLA2G15 in the presence of 20 µM amiodarone and fosinopril compared to no inhibitor control. Data are mean ± SD of *n* = 3 independent replicates. ****p < 0.0001. See also Extended Data Fig. 1-3.

## RESULTS

### PLA2G15 hydrolyzes BMP lipids

To test if BMP lipids are susceptible to phospholipase A (PLA)-mediated hydrolysis, we first incubated brain and liver lysates with commercially available BMP with the physiologically relevant stereoconfiguration (*S,S* BMP) ^1^ and monitored the release of the LPG intermediate (Fig. 1d). Consistent with a previous report ^9^, we observed a BMP hydrolase activity under both acidic and alkaline conditions, with relatively higher activity at neutral and mildly alkaline pH (Fig. 1d), probably due to the abundance of extralysosomal PLAs in the lysate with minimal lysosomal protein contribution ^17,18^. While these results indicate that BMP is not inherently resistant to PLAs, BMP could be protected by residing in the lysosome^19,20^. To test if lysosomal PLAs can degrade BMP lipids under acidic pH, we purified lysosomes from both tissues ^10,21^. Indeed, lysosomal lysates efficiently hydrolyzed BMP with a pH optimum of 4-5, reminiscent of that of the acid PLA (Fig. 1e) ^10^. Of importance, this activity was diminished by amiodarone, a non-specific inhibitor of lysosomal PLAs ^22,23^ (Fig. 1f). These data suggest that BMP lipids are vulnerable to degradation by lysosomal phospholipases^7,13–15^, indicating that their unique stereoconfiguration alone may not be sufficient to ensure their stability within the lysosome.

PLA2G15 is the major lysosomal phospholipase ^24–26^, and we recently established its function as a phospholipase B capable of catalyzing the complete hydrolysis of phospholipids with high efficiency ^10^. Given our results, we asked if PLA2G15 can degrade BMP lipids. To this end, we incubated lysosomal lipid extracts isolated from LysoTag mouse tissues ^10,21^ with purified PLA2G15 (Extended Data Fig. 1a-c) and monitored the abundance of glycerophospholipids with a focus on PG, the structural isomer of BMP (Fig. 1g). Consistent with its function, the abundance of most measured glycerophospholipids was significantly decreased (Extended Data Fig. 2a), with a concomitant increase in lysophospholipid intermediates (Extended Data Fig. 2b). Interestingly, most BMP lipid species were also significantly hydrolyzed independent of their acyl chain length and saturation (Fig. 1h). Of importance, no change was observed in the levels of sphingomyelin and triglycerides ^25,26^ (Extended Data Fig. 2c), suggesting a specific phospholipase activity towards BMP rather than an overall reduction in lipid abundance during the reaction. Similar results were obtained when using lysosomal lipids derived from HEK293T cells as substrates incubated with PLA2G15 (Extended Data Fig. 2d-g). Finally, using synthesized *S,S* BMP, the physiological stereoisomer, we validated the activity of PLA2G15 towards BMP using thin layer chromatography (Extended Data Fig. 2h) ^27^. Importantly, BMP hydrolysis was abolished by purified PLA2G15 S198A catalytic mutant (Extended Data Fig. 3a-c) ^10,28^ as well as in the presence of amiodarone or fosinopril, known inhibitors of phospholipid hydrolysis ^22^, thus confirming BMP hydrolase activity (Fig. 1i,j). Additionally, we observed no BMP hydrolase activity by purified PLBD2 (Extended Data Fig. 3d-f), an inositol-phospholipid preferring lysosomal phospholipase ^10^. Altogether, our data indicate that BMP lipids are susceptible to PLA2G15 activity.

### *sn2, sn2’* esterification position of *S,S* BMP confers resistance to PLA2G15 in vitro

While BMP levels significantly decreased after incubating lysosomal lipids with recombinant PLA2G15, a resistant fraction remained (Fig. 1h and Extended Data Fig. 2f). BMPs can be defined by acyl chain composition, position, and stereochemistry of chiral carbons (Fig. 2a). To gain mechanistic insight into their hydrolysis, we first performed kinetic studies of PLA2G15 activity towards BMPs with different acyl chain lengths under optimum conditions (Extended Data Fig. 4a-d) ^10,24^. Consistent with its role as a highly efficient phospholipase B ^10^, kinetic analyses of PLA2G15 monitoring the LPG intermediate and GPG product from the first and second steps, respectively, demonstrated high catalytic efficiencies (k_cat_/K_m_ = ∼10^5^ – 10^7^ M^−1^ s^−1^) against two commercially available BMP lipids, which were comparable to those against PG counterparts (Extended Data Fig. 4e-l) ^10^. As expected, the first step (Extended Data Fig. 4e-h) is faster than the overall reaction (Extended Data Fig. 4i-l), indicating that the second step controls the rate of the overall reaction ^10^. These results establish PLA2G15 as a potent hydrolase of BMP. While BMP usually displays *S,S* stereoconfiguration, other forms have been isolated including the *S,R* stereoisomer (Fig. 2a) ^4,29–32^. Therefore, we tested PLA2G15 activity against all three potential BMP stereoisomers with same acyl chains at the primary glycerol carbon positions (*R,R*, *S,R,* and *S,S*). To our surprise, all stereoisomers were equally hydrolyzed in short time experiments (30 seconds) monitoring LPG release and longtime ones (12 hours) monitoring remaining BMP levels (Fig. 2a,b and Extended Data Fig. 5a). Endogenous BMP can be acylated either on the secondary carbons *sn2, sn2’* (2,2’ BMP) or primary carbons *sn3, sn3’* (3,3’ BMP) of the glycerol (Fig. 2a) ^32,33^. Though the 3,3’ BMP is considered thermodynamically stable and readily synthesized, the 2,2’ BMP is thought to be a more abundant and physiologically active isoform ^32–36^. To examine BMP hydrolase activity against positional isomers, we obtained synthesized 2,2’ BMP and 3,3’ BMP standards, both of which contain a small amount of 2,3’ positional isomer that can be separated using our mass spectrometry methods and thus leveraged to monitor the stability of all positional isomers (Fig. 2a and Extended Data Fig. 5b). We found a striking reduction in BMP hydrolysis from 2,2’ BMP monitoring total BMP levels after prolonged incubation (12 hour), compared to 3,3’ BMP (Fig. 2b), despite only a milder difference observed when monitoring LPG release following short time (30 sec) incubation (Extended Data Fig. 5c). Intriguingly, a similar observation regarding the resistance of 2,2’ BMP is made when using lysosomal lysates (Fig. 2c). Of importance, these lysates are derived from cells deficient in CLN5, a known BMP synthase, and thus have very low levels of endogenous BMP ^2^. To further understand the hydrolysis of BMP isomers, we monitored time-dependent total BMP depletion for all isoforms incubated with PLA2G15 (Fig. 2d,e). Consistent with our earlier observation (Fig. 2b and Extended Data Fig. 5a), all three stereoisomers were hydrolyzed at the same rate (Fig. 2d). However, 2,2’ BMP has a much slower rate of hydrolysis compared to 3,3’ BMP when incubated with PLA2G15 (Fig. 2e), suggesting that esterification at the secondary carbon contributes to BMP’s resistance to hydrolysis. Consistent with this, we found the minimal reduction observed in the total BMP in the reaction containing 2,2’ BMP is mainly attributed to the hydrolysis of the 2,3’ positional isomer (Fig. 2f). Interestingly, we found that LPG released from 2,2’ BMP is stable and not degraded to GPG during time course experiments (Extended Data Fig. 5d,e), thus explaining why LPG monitoring gives a different result (Extended Data Fig. 5c) and establishing that monitoring LPG release could be misleading in studying the hydrolysis of BMP structural isomers. Additionally, when an equimolar mixture of both synthesized 2,2’ and 3,3’ BMPs is monitored for total BMP depletion, or LPG and GPG release, the results resemble the average of each positional isoform, supporting the notion that 3,3’ BMP is easily degraded compared to 2,2’ BMP (Fig. 2e and Extended Data Fig. 5d,e). Indeed, careful analysis of all positional BMP isoform peaks confirms that 3,3’ BMP is quickly degraded, followed by 2,3’ BMP, while the 2,2’ BMP is resistant (Fig. 2g,h).

**Fig. 2:**
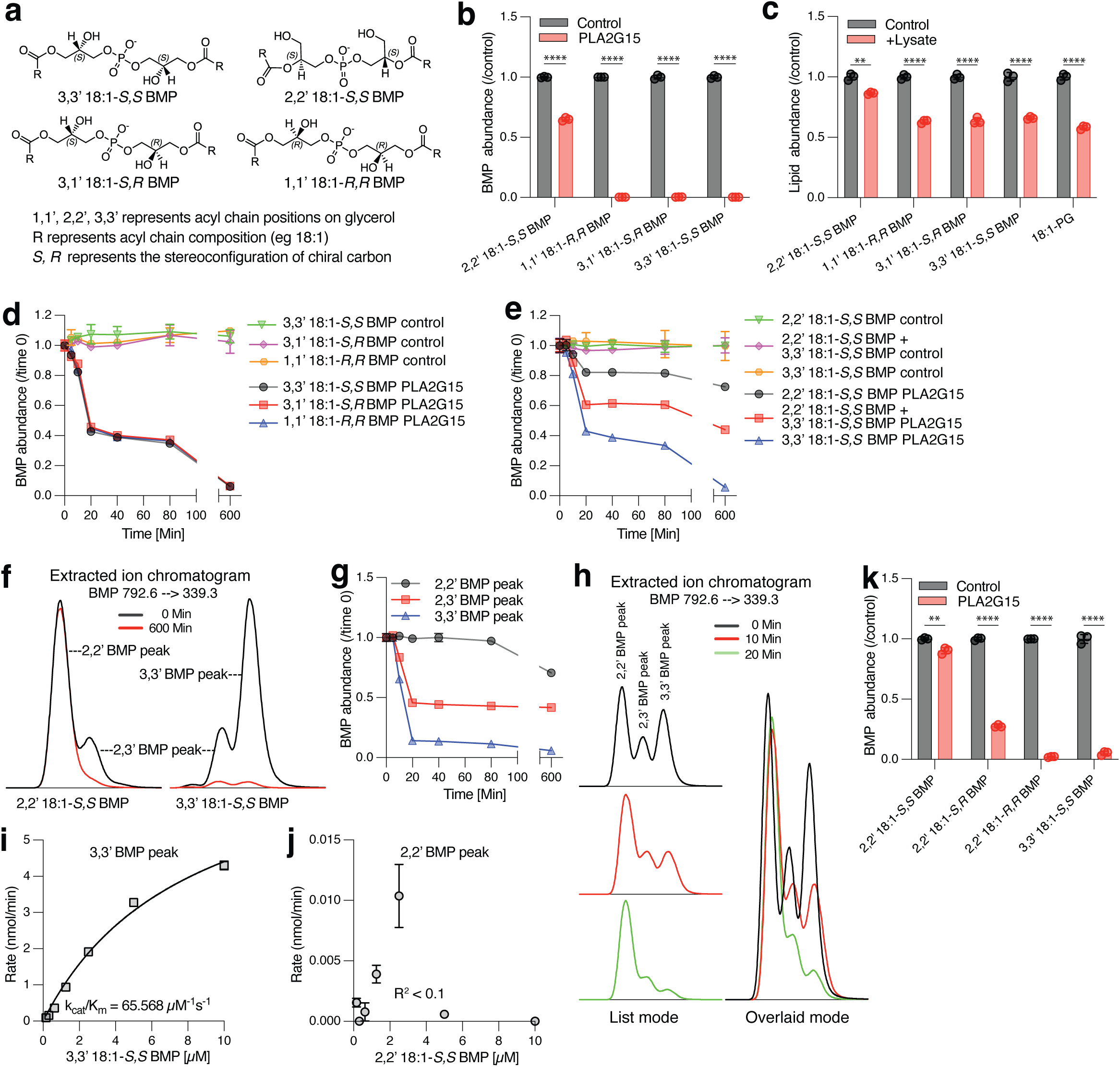
*sn2, sn2’* esterification position provides *S,S* BMP with resistance to PLA2G15. **(a)** Chemical structures of main BMP lipids used in this study. **(b)** PLA2G15 hydrolyzes BMP independent of stereoconfiguration while positional isomers are differentially hydrolyzed. Hydrolysis of 1 µM of indicated BMPs with different stereo and positional isomers by 100 nM recombinant PLA2G15 for 12 hours under acidic conditions (pH=5.0). Control has similar reaction components without the enzyme. BMP hydrolase activity was determined using LC-MS by calculating the abundance of total BMP remaining. Data are mean ± SD of *n* = 3 independent replicates. ****p < 0.0001. **(c)** Lysosomal lysates derived from *CLN5* KO HEK293T cells exhibit differential phospholipase activity towards positional isomers but not stereoisomers. Assay as in (b) for 2 hours using 10 µg of lysosomal lysate. *n* = 3 individual replicates for each BMP isomer or PG substrate. **p < 0.01 and ****p < 0.0001. **(d)** Time dependent hydrolysis of BMP stereoisomers by PLA2G15 confirms BMP degradation is independent of stereoconfiguration. Hydrolysis of 20 µM 3,3’ *S,S* BMP, 3,1’ *S,R* BMP or 1,1’ *R,R* BMP stereoisomers by 100 nM recombinant PLA2G15 over 10 hours under acidic conditions (pH=5.0). Controls had similar reaction buffers except with no enzyme. BMP hydrolase activity was determined using LC-MS by calculating the abundance of total BMP remaining at the indicated time points. Data are mean ± SD of *n* = 3 independent replicates. **(e-h)** Time dependent BMP hydrolysis shows 2,2’ *S,S* BMP positional isomer to be resistant to PLA2G15-mediated hydrolysis in vitro. **(e)** Same as in (d) using 20 µM BMP with either 2,2’ *S,S* BMP or 3,3’ *S,S* BMP positional isomer as well as a mixture of equimolar amounts. Data are mean ± SD of *n* = 3 independent replicates. **(f)** A representative image of the extracted ion chromatograms from (e) showing BMP peaks for each isomer at indicated time points in overlaid mode. **(g and h)** Time course analysis from the reaction with equimolar mixture of 2,2’ BMP and 3,3’ BMP in (e) shows 3,3’ peak is preferentially decreased followed by 2,3’ peak while 2,2’ peak is unchanged. Intensity of each BMP peak was carefully integrated at each time point in (g) and a representative image of the extracted ion chromatograms at indicated time points in (h) showing peaks representing the three positional isomers of BMP both in list and overlaid modes. **(i and j)** PLA2G15 deacylates 3,3’ BMP with faster kinetics than 2,2’ BMP. PLA2G15 was incubated with 3,3’ 18:1-*S,S* BMP in (i) and 2,2’ 18:1-*S,S* BMP in (j) under acidic conditions (pH= 5.0) for 10 min and reactions were stopped by heating before quantifying 2,2’ BMP or 3,3’ BMP peaks. Each experiment was repeated at least three times, and a representative graph is shown using 3,3’ and 2,2’ BMP peaks for analysis. K_cat_/K_m_ is a measure of kinetic efficiency. The phospholipids in these experiments were incorporated in liposomes with non-cleavable 18:1 Diether PC. **(k)** *S,S* stereochemistry in 2,2’ BMP is required for resistance against PLA2G15 hydrolysis. Assay as in (b) for 1 hour using 20 µM of indicated BMPs. Data are mean ± SD of *n* = 3 independent replicates. ****p < 0.0001. See also Extended Data Fig. 1,4,5.

Substrates bind to enzyme active site for effective catalysis and BMP is thought to activate lysosomal hydrolases including PLA2G15 ^3,37^. Interestingly, in silico docking experiments identified a plausible binding pocket that positioned the carbonyl carbon of BMP near the active site of PLA2G15 ^28^ (Extended Data Fig. 5f). Indeed, BMP binds inactive PLA2G15 (Extended Data Fig. 5g) indicating a similar phospholipase mechanism for BMP by PLA2G15 that involves the serine, aspartate, and histidine catalytic triad ^28^. Rigorous kinetic analyses monitoring individual peaks indicated that PLA2G15 has high catalytic efficiency against 3,3’ BMP peak (k_cat_/K_m_ = 10^7^ M^−1^ s^−1^) while there is no concentration dependent enzyme activity for 2,2’ BMP peak (Fig. 2i,j and Extended Data Fig. 4d). Finally, hydrolysis of BMP stereoisomers with *sn2, sn2’* acyl chain positions indicated that both esterification and stereochemistry (2,2’ *S,S*) are required for BMP resistance to degradation as 2,2’ BMPs with either *S,R* or *R,R* stereochemistry are readily hydrolyzed (Fig. 2k and Extended Data Fig. 5h,i). Altogether, these results elucidate PLA2G15 as a BMP hydrolase and demonstrate that the characteristic 2,2’ esterification position to be necessary to protect the physiological *S,S* BMP from hydrolysis.

### PLA2G15-deficient cells and tissues accumulate BMP

Several enzymes were reported to have catabolic activity against BMP in vitro, yet neither was validated as BMP hydrolase in cells or tissues ^7,9,13–16^. Targeted lipidomics revealed a significant increase in almost all BMPs in PLA2G15-deficient HEK293T ^10^ lysosomes and cells compared to their wildtype (WT) counterparts (Extended Data Fig. 6a,b and Supplementary Table 1). These results were further confirmed in additional clones (Fig. 3a,b, Extended Data Fig. 6c and Supplementary Table 2). Using our established lysosomal enzyme supplementation protocol ^2^, we delivered WT or catalytically dead PLA2G15 proteins (Extended Data Fig. 6d,e) to PLA2G15-deficient lysosomes and confirmed that these increases are dependent on the loss of PLA2G15 catalytic activity (Fig. 3c). Interestingly, the levels of one fifth of BMP were not rescued by PLA2G15, hinting that either longer treatment is required or that there may be other enzymes contributing to BMP degradation (Extended Data Fig. 6f) ^7,9,13–16^. Of note, many of the non-degraded BMPs have at least one 22:6 acyl chain, which showed a mixed phenotype in the *PLA2G15* knockout clones (Fig. 3a and Supplementary Table 2), indicating an interesting biology that is yet to be discovered ^11,29,32,34^. A recent paper suggested that PLA2G15 contributes to BMP synthesis in HeLa cells though changes observed were very minimal if any and the authors used a knockdown strategy ^38^. Using CRISPR knockout, we find significant increase in most BMPs of PLA2G15-deficient HeLa cells while 22:6-containing BMPs again showed a mixed phenotype (Extended Data Fig. 6g and Supplementary Table 3). Consistent with our cellular data (Fig. 3a,b), there is a significant increase in most BMPs isolated from brain, kidney, and liver of PLA2G15-deficient mice compared to their wildtype counterparts (Fig. 3d-f and Supplementary Table 4). Of importance, sphingomyelin, a non-glycerophospholipid species, remained mostly unchanged (Fig. 3g and Extended Data Fig. 6h) as previously reported ^25,26^. Despite PLA2G15 function as a phospholipase, there were minimal phospholipid changes if any, in PLA2G15-deficient lysosomes, cells and even tissues and changes in Hemi-BMP or acyl-PG levels were mixed and insignificant (Extended Data Fig. 6h and Supplementary Table 5). Finally, the levels of CLN5, a BMP synthase, and other lysosomal markers did not increase and so is the BMP synthesis in PLA2G15-deficient cells (Extended Data Fig. 6i,j). Thus, PLA2G15 loss of function increases BMP levels by reducing its degradation, indicating a vital role for PLA2G15 in BMP hydrolysis and homeostasis.

**Fig. 3:**
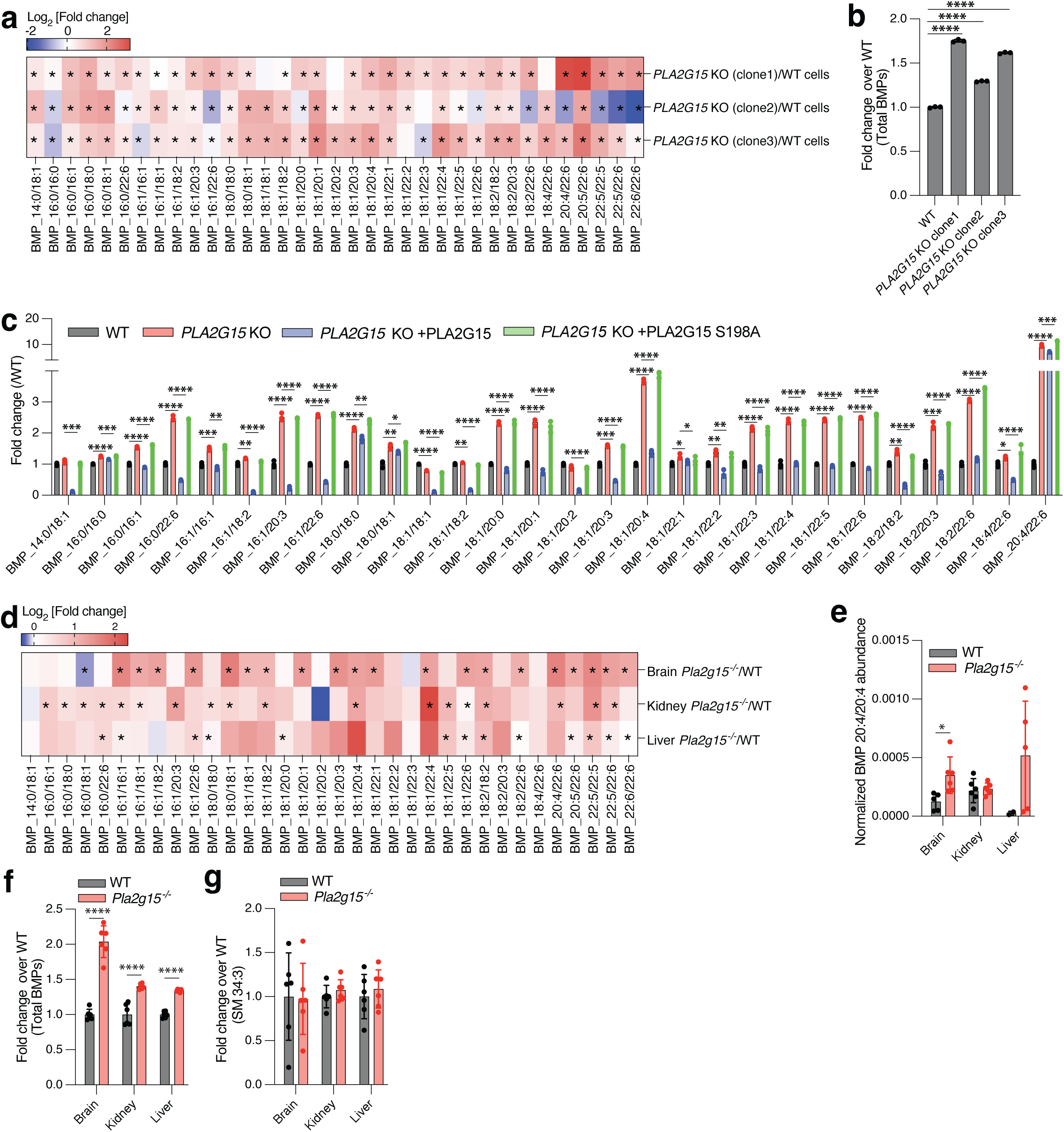
PLA2G15-deficient cells and tissues accumulate BMP. **(a and b)** Targeted analyses of BMP lipids reveal that a deficiency in PLA2G15 increases BMP levels in HEK293T cells. (a) Heatmap representation of log2-transformed changes in BMP abundances in PLA2G15-deficient HEK293T cells compared to wildtype controls (*n* = 3 WT and *n* = 3 *PLA2G15* KO). Data are the ratio of the mean of each BMP. Significant changes from the graph are represented by *p < 0.05. P values were calculated using two-tailed unpaired tests and are presented in Supplementary Table 2. (b) Fold changes in total BMP abundances. The sum of all abundances of measured BMPs was used to generate these values. Data are mean ± SD. Statistical analysis was performed using two-tailed unpaired *t*-tests. ****p < 0.0001. **(c)** Recombinant PLA2G15 rescues elevated levels of most BMPs resulting from PLA2G15 loss. Fold changes in the levels of BMPs in *PLA2G15* KO HEK293T cells after supplementation with wildtype or mutant PLA2G15 (*n* = 3 WT, *n* = 3 *PLA2G15* KO, *n* = 3 *PLA2G15* KO +PLA2G15 and *n* = 3 *PLA2G15* KO +PLA2G15 S198A). Data are mean ± SD. Statistical analysis was performed using two-tailed unpaired *t*-tests. *p < 0.05, **p < 0.01, ***p < 0.001 and ****p < 0.0001. **(d-f)** Targeted analyses reveal the accumulation of BMP lipids in mouse tissues with PLA2G15 deficiency. **(d)** Heatmap representation of log2-transformed changes in BMP abundances in PLA2G15-deficient mouse tissues (brain, kidney and liver) compared to their control counterparts (*n* = 6 WT and *n* = 6 *Pla2g15*^−/−^). Data are the ratio of the mean of each BMP. Significant changes from the graph are represented by *p < 0.05. P values were calculated using two-tailed unpaired tests and are presented in Supplementary Table 4. **(e)** Normalized abundances of BMP 20:4/20:4 in the same samples. This lipid was undetected in some samples; thus, it is reported separately. Data are mean ± SD. Statistical analysis was performed using two-tailed unpaired *t*-tests. *p < 0.05. **(f)** Fold changes in total BMP abundances. The sum of all abundances of measured BMPs was used to generate these values. Data are mean ± SD. Statistical analysis was performed using two-tailed unpaired *t*-tests. ****p < 0.0001. **(g)** Non PLA2G15 substrates are unchanged in PLA2G15-deficient mouse tissues. Fold changes in sphingomyelin (SM 34:3) in the same samples. Data are mean ± SD. See also Extended Data Fig. 6.

### PLA2G15 regulates lysosomal lipid metabolism through BMP

To further validate the role of PLA2G15 activity in regulating BMP homeostasis in the lysosome and the impact of this role on lysosomal function, we first determined that lysosomal lysates from PLA2G15-deficient HEK293T cells indeed have lower degradative activity against BMP (Fig. 4a). Next, we traced the fate of *S,S* BMP lipids in the presence and absence of PLA2G15 in cells by delivering BMP lipids to lysosomes using established protocols (Fig. 4b) ^2,10,39^. Of importance, we knocked out *PLA2G15* (Extended Data Fig. 7a) in *CLN5* knockout cells, whose endogenous BMP levels are minimal thus allowing tracing of unlabeled exogenously-delivered BMP (Extended Data Fig. 7b) ^2^. Of note, LC-MS analysis after delivery of either 2,2’ BMP or 3,3’ BMP to HEK293T cells indicated 2,2’ BMP as the major isoform in cells (Extended Data Fig. 7c), as previously reported ^32,34^. Using a pulse-chase approach to track BMP hydrolysis with or without PLA2G15 protein supplementation (Extended Data Fig. 7d,e) following exogenous 2,2’ BMP delivery to *CLN5 PLA2G15* double knockout cells (Fig. 4b and Extended Data Fig. 7f), we found that BMP is hydrolyzed faster in lysosomes with catalytically active PLA2G15 (Fig. 4c and Extended Data Fig. 7g). To determine the effect of the esterification position, we delivered either 2,2’ BMP or 3,3’ BMP and monitored their hydrolysis over time (Fig. 4b,d and Extended Data Fig. 7h,i). Careful peak analysis for 4-hour chase following 2,2’ BMP delivery showed a moderate 2,2’ peak decrease with a concomitant time-dependent increase in both 2,3’ and 3,3’ peaks (Fig. 4d and Extended Data Fig. 7h), suggesting the need for a conversion of 2,2’ BMP to 3,3’ BMP for effective degradation in cells. Consistent with this notion and our in vitro data (Fig. 2 and Extended Data Fig. 5), 3,3’ and 2,3’ peaks from 3,3’ BMP rapidly decreased especially in the presence of PLA2G15 while conversion of 3,3’ BMP to 2,2’ BMP is minimal if any (Fig. 4d and Extended Data Fig. 7h). To confirm this observation and minimize positional isomer conversion, we delivered an equimolar mixture of 2,2’ BMP and 3,3’ BMP (Fig. 4b,e and Extended Data Fig. 7J). Consistently, we observed rapid degradation of 3,3’ and 2,3’ BMP compared to 2,2’ BMP, confirming 2,2’ BMP as a more resistant BMP isomer (Fig. 4e,f). Notably, 2,2’ BMP peak is not affected by active PLA2G15 (Fig. 4e), suggesting that acyl migration is required for degradation by PLA2G15 (Fig. 4e,f). Importantly, *S,S* stereoconfiguration with primary esters (3,3’) does not protect BMP from hydrolysis in cells (Extended Data Fig. 7k).

**Fig. 4:**
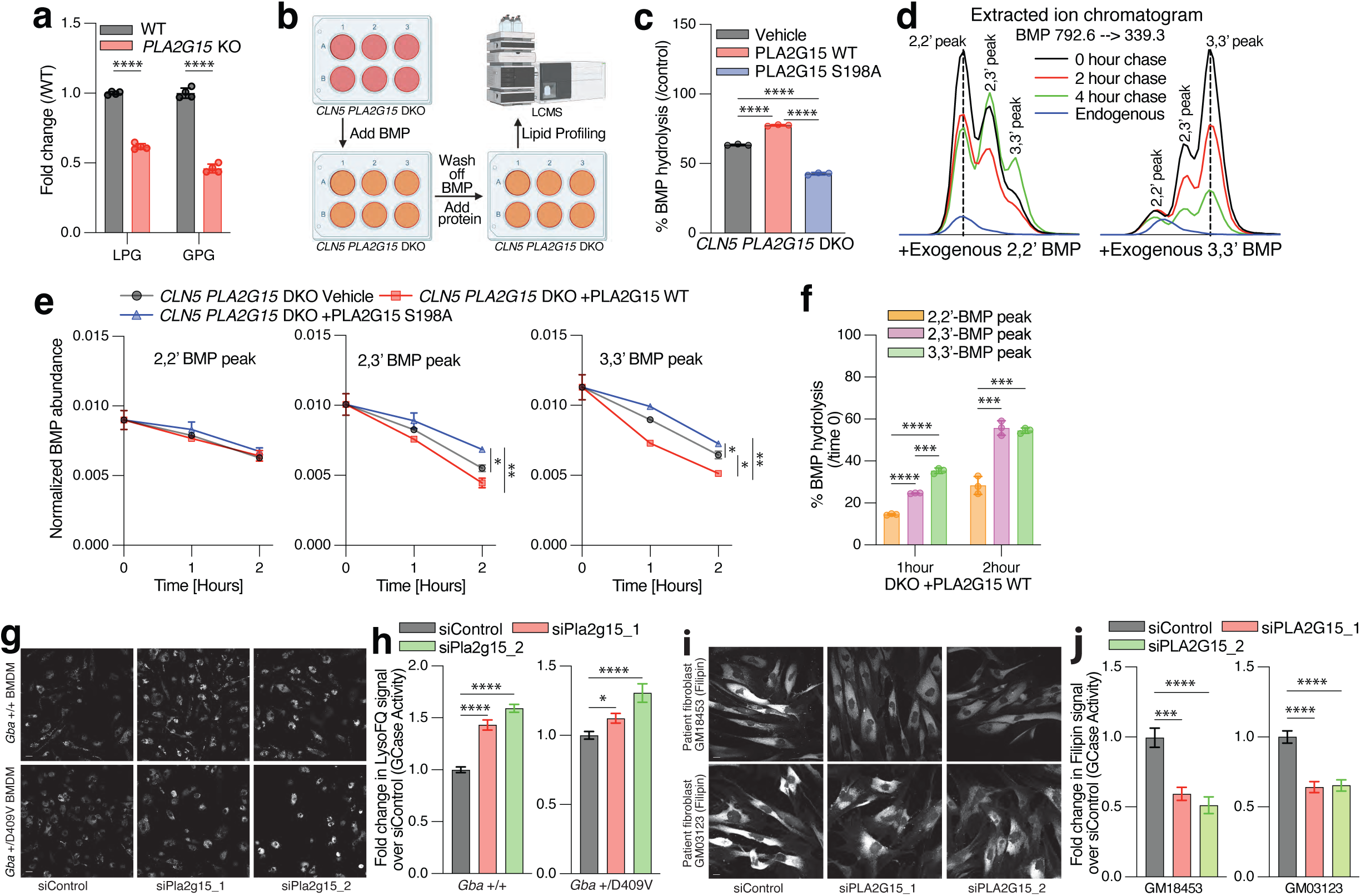
PLA2G15 regulates lysosomal lipid metabolism through BMP. **(a)** Lysosomes derived from *PLA2G15* KO cells exhibit reduced BMP hydrolase activity. Hydrolysis of 1 µM 3,3’ *S,S* BMP by 5 µg WT or *PLA2G15* KO lysosomal lysates for 10 min under acidic conditions (pH=5.0). BMP hydrolase activity was determined using LC-MS by measuring the abundance of LPG intermediate and GPG product. Data are mean ± SD. *n* = 3 individual replicates for each genotype. P values were calculated between *PLA2G15* KO versus WT. ****, p < 0.0001. **(b)** Depiction of experimental design for pulse-chase experiments. Times varies depending on the experiment. **(c)** PLA2G15 hydrolyzes BMP in cells, while inactive PLA2G15 S198A disrupts cells’ ability to degrade BMP. 10 µM 2,2’ *S,S* BMP was added for two hours to cells and then washed, and 100 nM protein was supplemented to cells. Measurements were done by LC-MS after 2-day chase. PBS was used as vehicle. Data are mean ± SD. *n* = 3 individual replicates for each condition. Statistical analysis was performed using two-tailed unpaired *t*-tests. ****p < 0.0001. **(d)** PLA2G15 degrades 3,3’ and 2,3’ *S,S* BMP from 3,3 *S,S* BMP while 2,2’ *S,S* BMP is converted to 2,3’ and 3,3’ BMP. Representative images of the extracted ion chromatograms showing all three positional BMP isomers peaks during a 4-hour PLA2G15 WT chase after delivering 2,2’ BMP (left) and 3,3’ BMP (right). **(e and f)** PLA2G15 preferentially degrades 3,3’ *S,S* BMP while 2,2’ *S,S* BMP turnover is slower. Time 0 represents the time before proteins were added after BMP wash off. **(e)** 2,2’ BMP peak (first), 2,3’ BMP peak (second) and 3,3’ BMP peak (third) was measured following a 2-hour chase with PLA2G15 proteins after addition of equimolar mixture of 2,2’ *S,S* BMP and 3,3’ *S,S* BMP for 2 hours. P values were calculated by one-way ANOVA. *p < 0.05, **p < 0.01. **(f)** Esterification position determines the rate of BMP hydrolysis. Quantitation of BMP peaks from (e) during the chase time with PLA2G15 WT confirms 3,3’ and 2,3’ BMP isomers are readily hydrolyzed while 2,2’ BMP may require conversion for efficient degradation. **(g and h)** PLA2G15 knockdown increases GCase activity in bone marrow derived macrophages (BMDMs) derived from wildtype and *Gba* heterozygous mutant mice. **(g)** Representative fluorescence microscopy images of BMDMs following a 30-minute incubation of 10 µM LysoFQ-GBA after three-day treatment with 10 nM control siRNA or two different siRNAs that target PLA2G15. Scale bar is 20 µm. **(h)** Quantification of signal intensity of LysoFQ-GBA. Signals are normalized to the siControl-treated BMDMs. *, p < 0.05, ****, p < 0.0001, by one-way ANOVA. **(i and j)** PLA2G15 knockdown reduces cholesterol phenotype in NPC1 patient-derived fibroblasts. **(i)** Representative microscopy images of Filipin staining of two independent NPC1 patient fibroblast cell lines (Coriell GM03123 and GM18453) after two-day treatment with 10 nM control siRNA or two different siRNAs that target PLA2G15. Scale bar is 20 µm **(j)** Quantification of signal intensity of Filipin. Signals are normalized to the siControl-treated patient fibroblasts. ****, p < 0.0001, by one-way ANOVA. See also Extended Data Fig. 7-9.

BMP is crucial for lysosomal lipid metabolism, in particular glucocerebrosidase (GCase) activity and cholesterol efflux ^2,12,20,36,40–42^. Consistent with PLA2G15 function as BMP hydrolase, we found that knocking down PLA2G15 upregulates GCase activity in bone marrow-derived macrophages (BMDMs) isolated from wildtype and heterozygous mutant *Gba* D409V mouse while supplementation of the active PLA2G15 decreases GCase activity in BMDMs (Fig. 4g,h and Extended Data Fig. 8a-c). BMP supplementation rescues cholesterol phenotype through an NPC2-dependent process in NPC1-deficient cells ^42–44^. Thus, we asked whether PLA2G15 inhibition can ameliorate cholesterol phenotype in NPC1 disease models. Consistent with this hypothesis, PLA2G15 was identified as a genetic modifier for NPC1 using a genome-wide haploid genetic screen using cholesterol staining with Perfringolysin O (PFO), a bacterial toxin that binds to cholesterol rich membranes and accumulates in NPC1-deficient cells (Extended Data Fig. 9a-f and Supplementary Table 6). Importantly, a few lysosomal enzymes are differentially regulating the cholesterol phenotype in NPC1-deficient cells (Extended Data Fig. 9d,e). In addition, the screen revealed several genes implicated in cholesterol uptake and efflux as strong regulators while other implicated pathways included glycosylation, organelle contact sites, cargo sorting, clathrin-mediated endocytosis, receptor trafficking/recycling, endosome maturation/trafficking and mitochondrial function (Extended Data Fig. 9a-f and Supplementary Table 6) ^45–47^. Consistent with the screen, RNAi-mediated knock down of PLA2G15 reduced cholesterol phenotype in two independent NPC1 patient fibroblast lines using Filipin stain as an orthogonal approach to detect cholesterol (Fig. 4i,j and Extended Data Fig. 8d). These data indicate that PLA2G15 regulates lysosomal lipid metabolism through BMP and its inhibition can ameliorate disease-related lysosomal lipid dyshomeostasis.

### Genetic targeting of *Pla2g15* reverses defects in NPC1-deficient mouse

To test if PLA2G15 depletion is of therapeutic value in NPC1 disease, we generated PLA2G15-deficient mice and crossed these to the *Npc1*^m1N^/J mouse, a model that exhibits severe central nervous system (CNS) and peripheral NPC characteristics (Fig. 5 and Extended Data Fig. 10). First, we measured well-established disease biomarkers including neurofilament light chain (Nfl) in cerebrospinal fluid (CSF) and plasma for neurodegeneration and aspartate aminotransferase (AST) and alanine aminotransferase (ALT) for liver damage ^48–50^. Depletion of PLA2G15 in NPC1-deficient mice significantly reversed these disease biomarkers (Fig. 5a,b). NPC1 models display increased BMP levels ^51^, thus PLA2G15 depletion shows only slight increase in a few BMP species if any, consistent with potential cell-type specific effect ^41^, or a net effect of reduced lysosomal stress in NPC1 lysosomes in the double knockout mice (Extended Data Fig. 10a and Supplementary Table 7). While not reaching significance, PLA2G15 loss of function slightly decreased the cholesterol phenotype detected in liver, however, elevated secondary storage lipids, including sphingolipids and alkyl-lysophosphatidylcholine, were reduced significantly in brain and liver (Extended Data Fig. 10b-d and Supplementary Table 8) ^52–54^. NPC1 deficiency profoundly affects Purkinje neuron survival and development, triggers glial reactivity and impacts multiple organ systems ^55–60^. Remarkably, depletion of PLA2G15 in NPC1-deficient mice significantly alleviated neuropathology findings such as Purkinje cell loss, astrocytosis, microgliosis and demyelination across the central nervous system (CNS) (Fig. 5c-f, Extended Data Fig. 10e,f and Supplementary Table 9). Also, inhibition of PLA2G15 lessened hyperplasia in Kupffer cells from NPC1-deficient liver, while vacuolation in hepatocytes remained unaltered (Extended Data Fig. 10g,h). Similarly, inhibition of PLA2G15 reduced the hyperplasia of histocytes and lymphoid atrophy in NPC1-deficient spleen, while histological findings observed in the lungs were not corrected (Extended Data Fig. 10g,h). Notably, histopathology and histomorphometry evaluations of the CNS, liver, spleen and lungs revealed no lesions in PLA2G15-deficient mouse compared to control mouse (Fig. 5c-f, and Extended Data Fig. 10g,h). Consistently, genetically targeting PLA2G15 strongly improved neurological composite score and ataxia symptoms in NPC1-deficient mice, leading to a significantly extended lifespan of disease mice (Fig. 5g-i and Extended Data Fig. 10i) ^55^. These data strongly suggest that PLA2G15 depletion reduces neurodegeneration in brain and defects in visceral organs of NPC1-deficient mice. Overall, these results, in conjunction with our in vitro and in vivo data, indicate that PLA2G15 possesses a BMP hydrolase activity that is required for BMP homeostasis in the lysosome, and its targeting can boost BMP-mediated functions and lysosomal activity in diseases with lysosomal dysfunction.

**Fig. 5:**
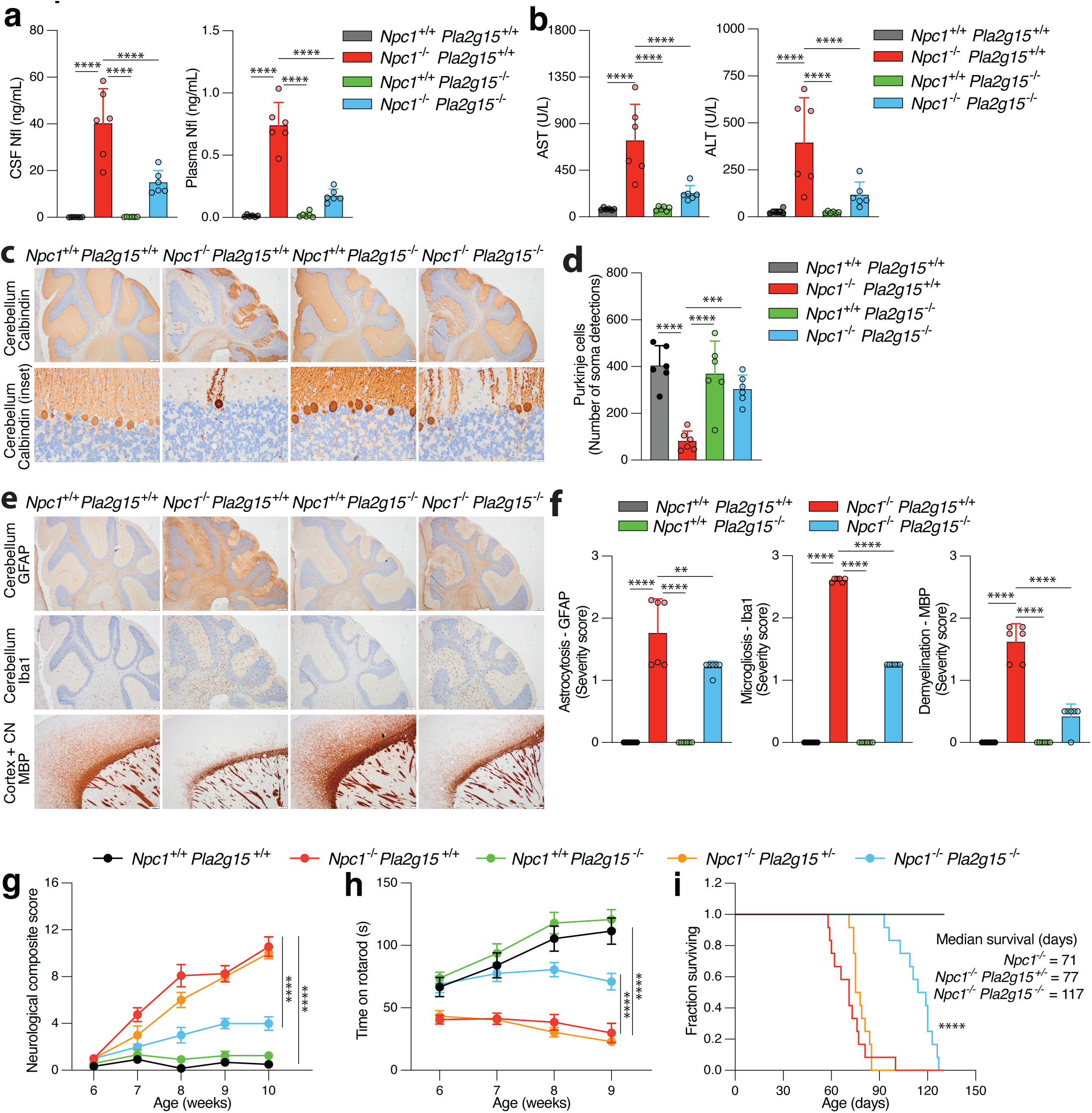
Genetic ablation of *Pla2g15* ameliorates disease symptoms and extends lifespan in NPC1-deficient mouse. **(a-i)** Genetic depletion of PLA2G15 in NPC1-deficient mouse. **(a and b)** PLA2G15 depletion rescues elevated levels of neurodegenerative and liver damage biomarkers in NPC1-deficient mice. Neurofilament light chain (Nfl) concentrations in CSF (left) and plasma (right) were measured in (a), while plasma levels of liver transaminases AST (left) and ALT (right) were measured in (b) at day 56. Data are mean ± SD (*n* = 6, 3 males and 3 females per genotype). *, p < 0.05, **, p < 0.01, ***, p < 0.001, ****, p < 0.0001, by one-way ANOVA. **(c-f)** Genetic depletion of PLA2G15 increases Purkinje cell survival (c and d) and decreases astrocytosis, microgliosis and demyelination (e and f) in the CNS of NPC1-deficient mice. Representative images are displayed in c and e. CN represents cerebral nuclei. Scale bars are 200 µm for all images except 20 µm for Cerebellum Calbindin inset. Histomorphometry to quantify the number of Purkinje cells per slide and the mean severity score of the histopathology evaluation in the CNS is shown in d and f respectively. Severity score is the mean grade of the neuropathology evaluation performed independently in the following neuroanatomical locations: cerebellum, hindbrain (pons and medulla), midbrain, interbrain (thalamus and hypothalamus), basal ganglia (striatum and pallidum), hippocampus, isocortex (somatomotor, somatosensory and visual areas) and olfactory bulb. Data are mean ± SD (n = 6, 3 males and 3 females per genotype) and **, p < 0.01, ***, p < 0.001, ****, p < 0.0001, by one-way ANOVA comparing NPC1-deficient tissues to other genotypes. **(g-i)** Genetic depletion of PLA2G15 improves neurological composite score (g), motor defect (h) and increases survival rate (i) in NPC1-deficient mice. Note: Higher neurological composite score means worse neurological performance and was calculated using hindlimb clasping, grooming, tremor, gait, kyphosis, and ledge test. Data are mean ± SEM (*n* = 12, 6 males and 6 females per genotype). In (h) ataxia symptoms were measured using rotarod. Data are mean ± SEM (*n* = 12, 6 males and 6 females per genotype). ****, p < 0.0001, by two-way ANOVA comparing *Npc1*^−/−^ *Pla2g15^+/+^* to either *Npc1*^−/−^ *Pla2g15^−/−^* or *Npc1*^+/+^ *Pla2g15^+/+^* **(i)** Kaplan-Meier graph for animal survival. Data from *n* = 12, 6 males and 6 females per genotype. The curves for *Npc1*^+/+^ *Pla2g15*^+/+^ and *Npc1^+/+^ Pla2g15*^−/−^ completely overlap. ****, p < 0.0001, by Log-rank (Mantel-Cox) test. See also Extended Data Fig. 10.

## DISCUSSION

BMP is a unique lipid known to potently stimulate lysosome-mediated lipid turnover, a crucial cellular process that, when defective, leads to a plethora of human diseases including neurodegeneration ^1,4^. Despite the therapeutic potential in targeting the BMP pathway and its significant functions in the lysosome, little is known about BMP synthesis and degradation ^1,4^. We have recently discovered CLN5 as a BMP synthase in the lysosome, however, the rules that govern BMP stability and the enzymes that mediate its hydrolysis, if any, are yet to be discovered. For over fifty years, the belief has been that lysosomes do not have BMP hydrolase or that the unique stereochemistry of BMP (*S,S*) alone is protective from such hydrolase if exists ^11,12^. Here, we identify the lysosomal phospholipase PLA2G15 as an efficient BMP hydrolase regardless of stereoconfiguration if BMP has primary esters, challenging both assumptions and leading us to ask what protects BMP from hydrolysis within the lysosomal environment. Interestingly, we found that the *sn2, sn2’* (2,2’ BMP) esterification position, another unique feature of BMP, along with *S,S* stereoconfiguration, to be what shields BMP from lysosomal degradation, facilitated in part by PLA2G15. Consistent with these biochemical findings, loss of PLA2G15 leads to accumulation of BMP lipids in cells and tissues while most other phospholipids were unchanged ^25,26,61–63^. These results indicate that acyl migration within the BMP molecule is an active process that allows regulated turnover of BMP in the lysosome. Indeed, we observed that 2,2’ BMP is converted to 2,3’ BMP and 3,3’ BMP, both of which are susceptible to PLA2G15 activity. While this conversion is favored thermodynamically, we still found 2,2’ BMP to be more abundant than 3,3’ BMP in the lysosome, consistent with the notion of it being the biologically active form ^32–36^. We speculate that BMP stability is maintained by an unidentified biological machinery that protects its 2,2’ BMP isoform from acyl migration ^1^. Future studies should investigate the preferential synthesis or preservation of 2,2’ BMP, as well as the enzymes or processes required for its conversion to the 3,3’ isoform during its catabolism.

Consistent with BMP’s role in promoting lysosomal functions such as lipid degradation and cholesterol trafficking ^1–4,43,64^, we showed that inhibiting and supplementing PLA2G15 to BMDMs upregulates and reduces GCase activity, respectively, while depleting PLA2G15 can reduce the cholesterol phenotype in patient-derived NPC1-deficient fibroblasts. Importantly, we found that genetic depletion of PLA2G15 ameliorates secondary and primary lipid storage, neurodegeneration, spleen and liver damage leading to improved neurological symptoms and survival of NPC1-deficient mouse model ^48–50,55–58^. These data suggest that inhibiting PLA2G15 represents a potential therapeutic strategy to boost BMP levels and treat multiple lysosome-related diseases, an observation consistent with the BMP therapeutic hypothesis that we recently proposed based on work done by many others ^1^.

In summary, our work identifies PLA2G15 as a key hydrolase actively degrading BMP in the lysosome and establishes the rules that protect BMP from lysosomal hydrolysis, thus resolving the longstanding mystery of how BMP’s unique characteristics mediate its functions. These insights pave the way for new therapeutic strategies targeting the BMP pathway in diseases where it is disrupted.

## MATERIALS and METHODS

### Chemicals and Antibodies

Antibodies and chemicals used in this study are listed with their details in tables below.

### Antibodies

| Antibody | Company | Catalogue number or | Dilution |
| --- | --- | --- | --- |

|  |  | **Research Resource Identifiers** | **factor** |
| --- | --- | --- | --- |
| His | Cell Signaling Technology | 2365/ RRID:AB_2115720 | 1:1000 |
| LAMP2 | Santa Cruz Biotechnology | SC-18822/ RRID:AB_626858 | 1:1000 |
| Vinculin | Santa Cruz Biotechnology | SC-73614 | 1:1000 |
| CLN5 | Abcam | Ab170899 | 1:1000 |
| CTSB | Cell Signaling Technology | CST-31718 | 1:1000 |
| Calbindin | Abcam | Ab229915 | 1:1000 |
| MBP | Abcam | Ab218011 | 1:1000 |
| Iba1 | Abcam | Ab178846 | 1:1000 |
| GFAP | Abcam | Ab7260 | 1:1000 |
| Mouse HRP | Cell Signaling Technology | 7076/ RRID:AB_330924 | 1:3000 |
| Rabbit HRP | Cell Signaling Technology | 7074/ RRID:AB_2099233 | 1:3000 |

### Chemicals and reagents

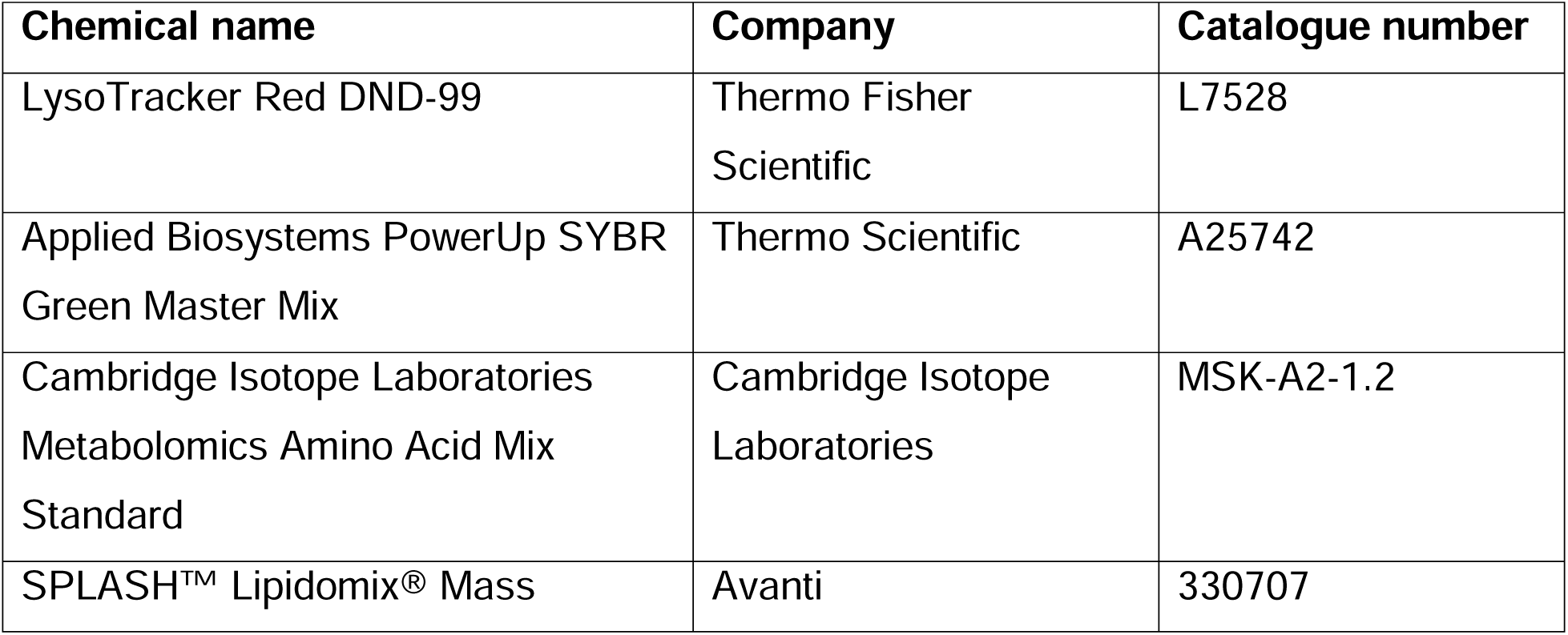
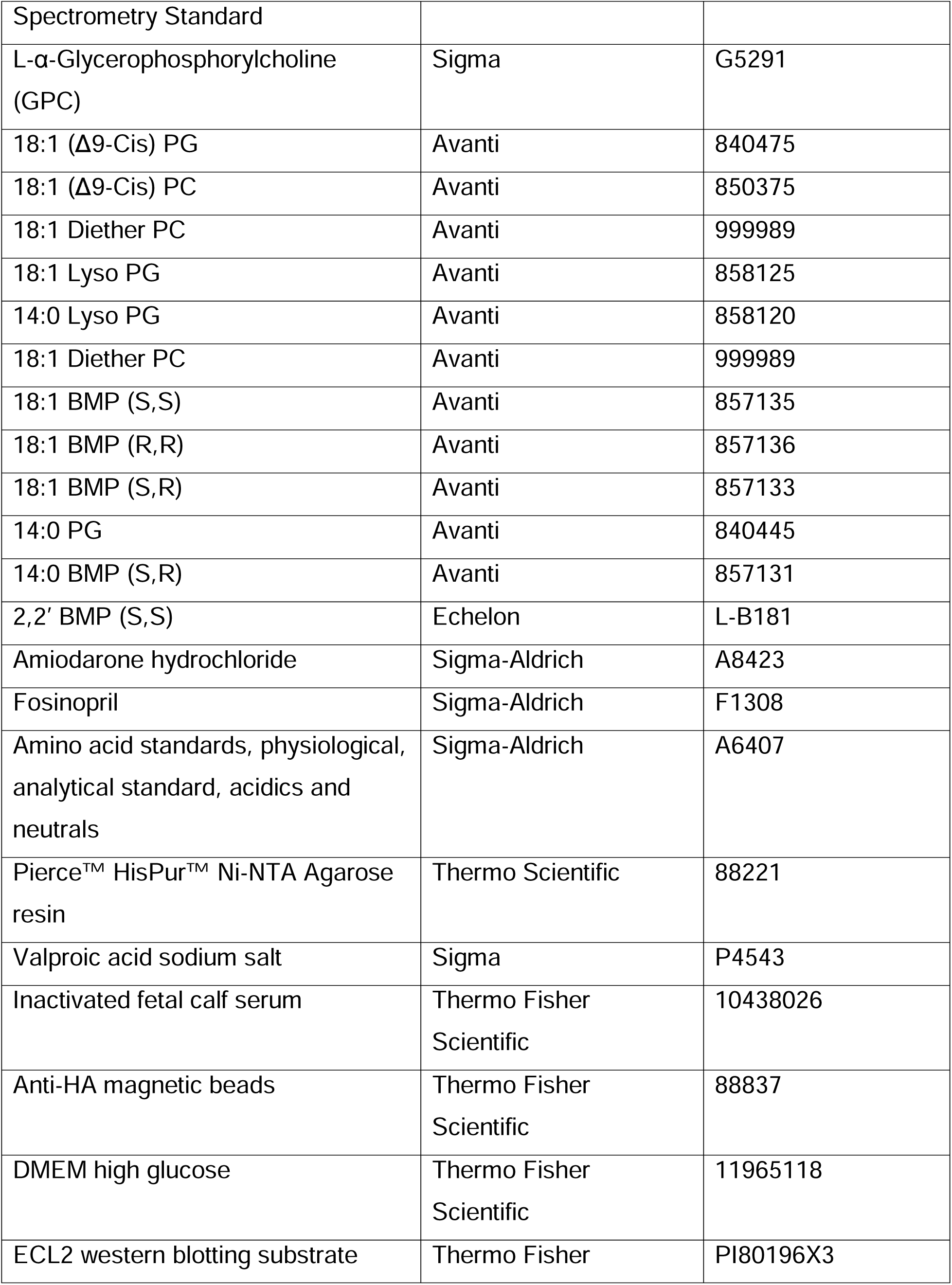
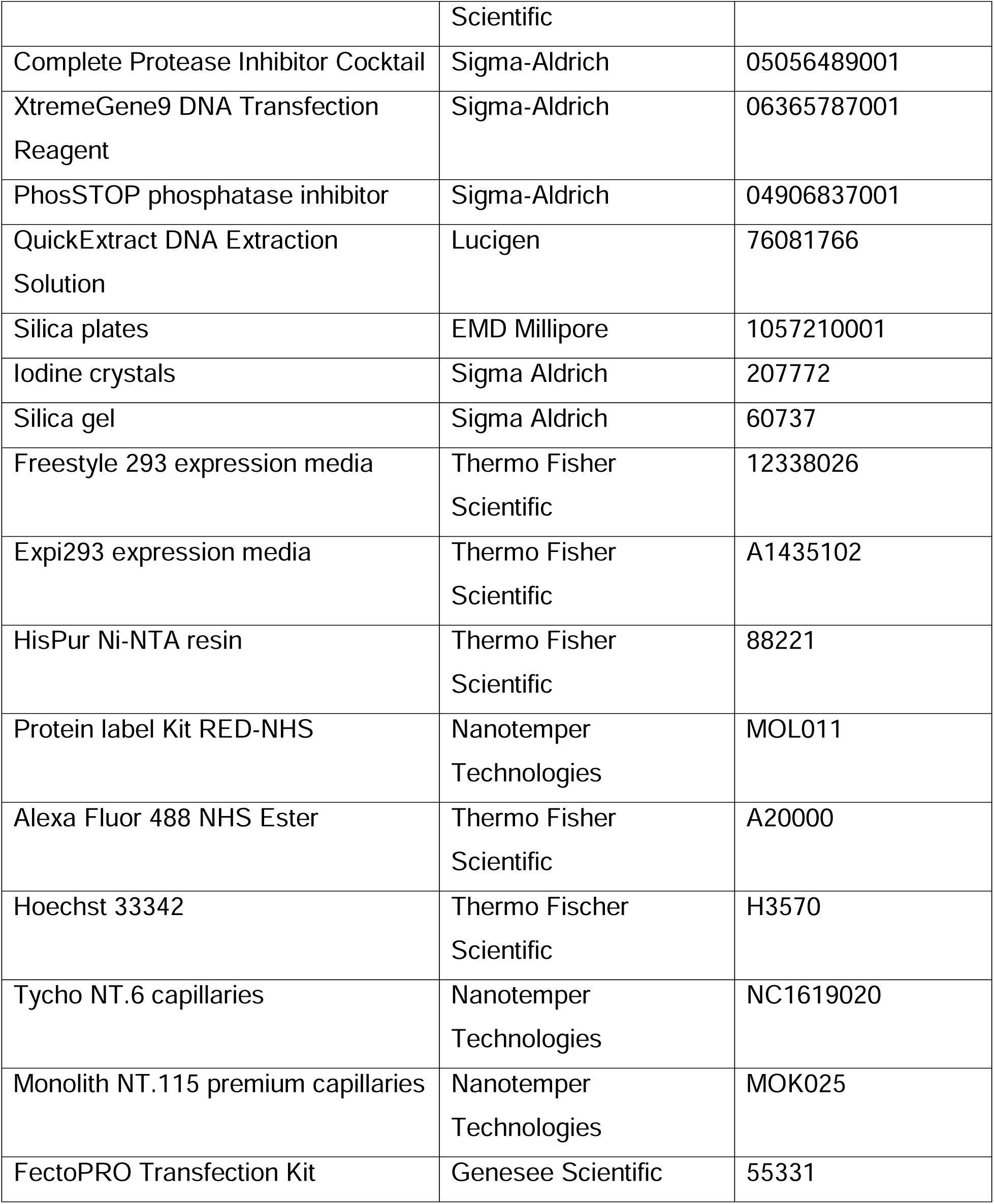

### Experimental models

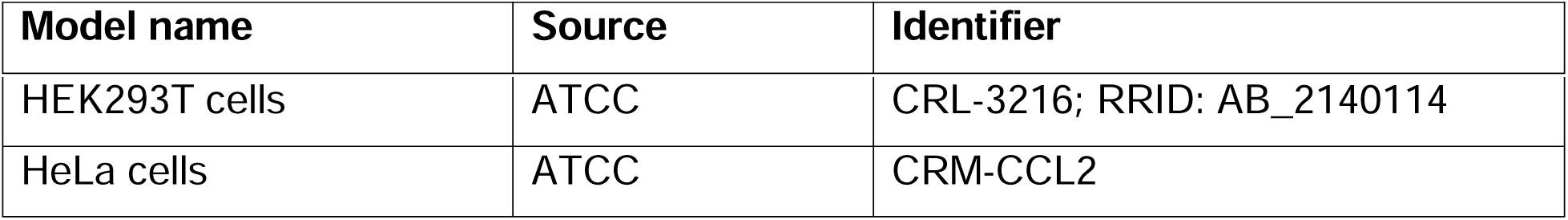

| Model name | Source | Identifier |
| --- | --- | --- |
| HEK293T cells | ATCC | CRL-3216; RRID: AB_2140114 |
| HeLa cells | ATCC | CRM-CCL2 |

### Software

| Software name | Source | Identifier |
| --- | --- | --- |
| Agilent LC/MS Data Acquisition software | Agilent | https://www.agilent.com |
| Agilent Qualitative Analysis software | Agilent | https://www.agilent.com |
| Agilent Qualitative Analysis software | Agilent | https://www.agilent.com |

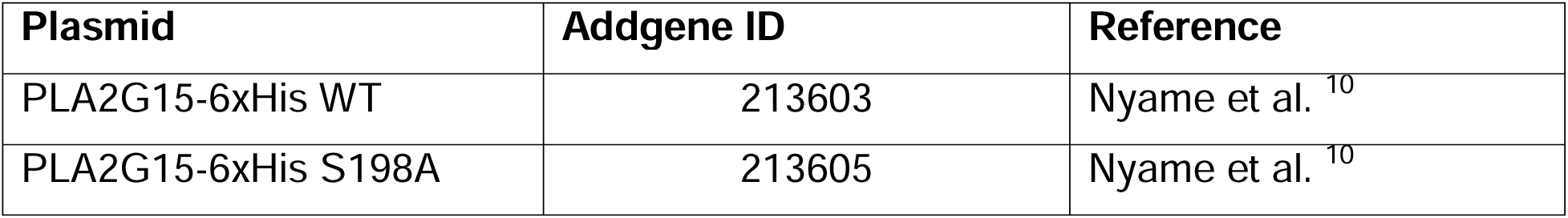

| Plasmid | Addgene ID | Reference |
| --- | --- | --- |
| PLA2G15-6xHis WT | 213603 | Nyame et al. <sup>10</sup> |
| PLA2G15-6xHis S198A | 213605 | Nyame et al. <sup>10</sup> |

### Plasmids

The Plasmids used in this study are in the plasmid table, and their references are included.

### Animal Studies

The mice were acquired and maintained as described before ^21^. The mice were housed in a controlled environment with regulated temperature and humidity, 12 hour light/dark cycles, and with access to food and water. Supplies were checked daily, and the cages were cleaned every 4-5 days. All mice procedures were conducted in adherence to the approved guidelines at Stanford University. LysoTag mouse was previously reported (Jax: Strain #:035401) ^21^. *Gba^D409V^* was obtained from Jax (strain # 019106) ^65^.

PLA2G15-deficient mice were obtained, housed, and used in accordance with approved IACUC protocols at the University of Wisconsin-Madison ^26,66^. Before euthanasia and necropsy, PLA2G15-deficient mice were fasted 6 hours prior.

Mouse genotypes were confirmed by PCR using DirectPCR (Thermo Fisher #AA4647500) according to manufacturer guidelines. Briefly, tail clippings were incubated in DirectPCR overnight at 60°C for 16 hours, proteinase K was added an incubated for 1 hour. PCR was performed with PLA2G14 primers (Pla2g15_1: 5’-GAATTCCTAGACCCCAGCAAGAGGAATGTG −3’; Pla2g15_2:5’-ACTGCTCCCCCTCCCCAGAGATGGATATTT-3’) using a program of 4 minutes at 95°C, followed by thirty-five cycles of 1 minute 95°C, 1 minute 55°C, 1 minute 72°C, and ended with an 8 minute incubation at 72°C. Results of genotyping were validated using RT-PCR and western blots of excised tissue. These mice were used in Fig. 3.

### Generation of *Pla2g15*^−/−^ mice for the NPC1 rescue experiment

Constitutive *Pla2g15*^−/−(TAC)^ mice were obtained via CRISPR/Cas9-mediated gene editing at Taconic Biosciences in Denmark in accordance with national and international regulations. Animal procedures were evaluated by veterinarians and Taconic’s IACUC or equivalent oversight body to ensure elimination or minimization of the potential for pain or distress. The gene targeting strategy was based on NCBI transcript NM_133792.3 (*Pla2g15*) and was designed to delete exon3 from the transcript (proximal sgRNA: 5’-TCGATGCCATTCATGGACTT-3’ and distal sgRNA: 5’-GGCCTACTACGTTGCTTTGA-3’), resulting in loss of function of PLA2G15 by generating a frameshift from exon 2 to all downstream exons and creating a premature stop codon in exon 4.

*Pla2g15* exon 3 flanking primers (forward: 5’-ACTCACTTTATAGACCAGGTTGGC-3’ and reverse: 5’-CAGAGACAGTGAACCTAAGGGC-3’) were designed to amplify and confirm both the wild-type and edited *Pla2g15* allele.

Briefly, Cas9 protein and sgRNAs were injected into BALB/c zygotes and G0 animals were genotyped by PCR analysis as described above. Founders were identified and used for further breeding activities to create heterozygous *Pla2g15*^−/−(TAC)^ mice. These mice were subsequently used for generation of homozygous *Pla2g15*^−/−(TAC)^ mice. These mice were used in experiments in Fig. 5.

### Life span analysis and neurological phenotyping

*Pla2g15*^−/−(TAC)^ Balb/C animals were crossed with *Npc1^m1N/J^* heterozygous (HET) animals (JAX #003092), and from the resulting offspring, double heterozygous mice were used to obtain the following genotypes (gene order: *Npc1^m1N/J^/Pla2g15*): WT/WT, HOM/WT, HOM/HET, HOM/KO and WT/KO. WT represents *Npc1*^+/+^ or *Pla2g15*^+/+^, KO represents *Npc1^+/+^ Pla2g15*^−/−^, HOM represents *Npc1*^−/−^ *Pla2g15^+/+^*, and HET represents *Pla2g15*^+/−^. In total 60 animals were enclosed for the phenotypic analysis, 12 animals of each of the different genotypes (6 male and 6 female). Bodyweight of all animals was assessed every other day starting at three weeks of age after the pups are weaned. Motor coordination was assessed weekly in the rotarod test from 6 to 9 weeks of age.

Neurological composite phenotype score consisting of ledge test, hindlimb clasping, gait, kyphosis, tremor, and grooming scores was performed on a weekly basis starting at 6 weeks of age. These tests were performed following established protocol ^67^. Higher composite score means worse neurological performance. Daily monitoring of clinical signs or termination criteria scoring and special care (e.g. wet food) exceeding standard health checks and animal care was assessed daily in all animals carrying the *Npc1^m1N/J^* HOM genotype starting at an age of 8 weeks). Experiments were performed at QPS Austria in compliance with the Animal Care and Welfare committee, as well as animal welfare regulations from local authorities.

### Cerebrospinal fluid and plasma analysis

At 8 weeks of age (P56 ± 2 days) animals were terminally anesthetized by intraperitoneal injection of Pentobarbital (600 mg/kg) and CSF and blood plasma was obtained. Neurofilament light chain (NfL) levels were assessed using ELISA 10-7001 CE from Uman Diagnostics in plasma and CSF. Plasma samples were diluted 1:3 and CSF 1:30 (wherever possible) in assay buffer and analyzed according to the manufacturers protocol.

AST and ALT levels were determined with a Kit (AST: Cat# 04467493190, Roche; ALT: Cat No 04467388190, Roche) according to International Federation of Clinical Chemistry and Laboratory Medicine (IFCC) with pyridoxal phosphate activation (Roche). Therefore, a kinetic measurement of the enzyme activity with a redox reaction of NADH was performed, using L-Aspartate and 2-Oxoglutarat as substrate for the AST measurement and L-Alanine and 2-Oxoglutarat as substrate for the ALT determination. The two enzyme levels were measured on a Roche Cobas 6000/c501 analyzer.

### Histoprocessing

The CNS, liver, spleen and lungs of all animals were sampled at necropsy, frozen and transferred to AnaPath Services GmbH for histoprocessing. Samples were placed in formalin, embedded in paraffin wax and cut at nominal thickness of 4µm. Liver, spleen and lungs were stained with hematoxylin and eosin (HE). In the CNS, five sagittal serial sections of the brain were performed: one section was stained with HE and four sections were employed for immunohistochemistry and labeled with antibodies against calbindin (ab229915; Lot Nummer_GR3361538-6; Abcam) for targeting Purkinje cells, myelin basic protein (MBP) (ab218011; Lot Nummer 1007654-30; Abcam) for targeting myelin sheaths, Iba1 (ab178846; Lot Nummer 1002201-50; Abcam) for targeting microglia and macrophages and glial fibrillary acidic protein (GFAP) (ab7260; Lot Nummer GR3454901-1; Abcam) for targeting astrocytes. Immunohistochemistry was performed with a BOND III (Leica Biosystems).

### Histopathology Evaluation

Histopathology evaluation was performed by an EBVS® European Veterinary Specialists in Pathology using an Olympus BX46 microscope. Illustrative microscopic images were taken with an Olympus SC50 camera. Evaluation of the CNS was performed independently in the following neuroanatomical locations: cerebellum, hindbrain (pons and medulla), midbrain, interbrain (thalamus and hypothalamus), basal ganglia (striatum and pallidum), hippocampus, isocortex (somatomotor, somatosensory and visual areas) and olfactory bulb. Histological changes were described according to distribution, severity, and morphologic character. Severity scores were assigned from a scale of 1-5 as follows: Grade 1, Minimal; Grade 2, Slight; Grade 3, Moderate; Grade 4, Marked; Grade 5, Severe. Histopathology findings recorded in different neuroanatomical locations of the CNS included: Purkinje cells loss (Decreased number of soma/decreased dendrites); Microgliosis (Increased number/activation of microglia); Astrocytosis (Increased number/activation of astrocytes); Demyelination; Foam cell formation. After the independent evaluation of the above-mentioned neuroanatomical regions, a mean severity score of each finding throughout the CNS was calculated. Histopathology findings in the other organs included: hyperplasia of Kupffer cells (Increased number/activation/foamy cytoplasm) and vacuolation of hepatocytes (Cytoplasmic alteration/glycogen/lipid accumulation) in the liver; Hyperplasia of histiocytes (Increased number/activation/foamy cytoplasm) and atrophy of lymphocytes (Decreased number in the while pulp) in the spleen; Alveolar macrophages (Activated/foamy cytoplasm) and cell debris in the alveolar lumen of the lungs.

### Histomorphometry

Histomorphometry analyses were performed on Calbindin-immunolabeled CNS sections in all the animals to quantify the number of Purkinje cells. Slides were scanned by an Olympus Slideview VS200 slide scanner coupled to an Olympus VS-264C camera using the 40x objective to obtain whole slide images (WSI). Histomorphometry evaluation was conducted in QuPath ^68^, quantitative pathology and bioimage analysis software, version 0.4.3. Cerebellum was annotated as the ROI (Region of Interest). Cell detection algorithm was employed for the detection of the soma of Calbindin-positive cells (i.e., Purkinje cells).

### Cell culture

HEK293T cells, HeLa cells and their derivatives were cultured in Dulbecco’s modified Eagle’s medium (DMEM; Gibco) base medium supplemented with 10% heat inactivated fetal bovine serum (FBS) (Thermo Fisher Scientific) with 2 mM glutamine, 100IU/mL penicillin, and 100µg/mL streptomycin (Thermo Fisher Scientific). Cell cultures were maintained at 37 °C in an incubator with 5% CO_2_. PLA2G15-deficient clone1 cells with LysoTag were obtained from the previous study ^10^. For protein expression, Expi293F cells were maintained in a shaking incubator at 37 °C with 8% CO_2_.

### Generation of knock-out (KO) cell lines using CRISPR-Cas9 technology

*PLA2G15* KO clone1 cells were described before ^10^. *PLA2G15* KO HEK293T clone 2, 3 and HeLa cells were generated following the Synthego Gene Knockout Kit v2 protocol and using the following mixed guide sgRNAs sequences:

GGGGCGGAGGUGGAGGCCCA
UUAGCAGCAGCAAGAGGAAC
CCUCACCCAGCACCACUGGG

In short, a combined 180 pmol sgRNA and 20 pmol Cas9 (IDT) were incubated at room temperature for 10 minutes. Concurrently, HEK293Ts were washed with PBS, detached using enzyme-free dissociation buffer (ThermoFisher), and counted to determine their density. A suspension of 150,000 cells was added to the newly formed RNP complexes, transferred to a Lonza Nucleocuvtte^TM^, and electroporated using the CM-130 program code. Following electroporation, cells were placed in growth medium and transferred to a 24-well plate. Single cell populations were obtained using limited dilution and the sgRNA target sites were sequenced following PCR-amplification. Finally, genetic knockouts were confirmed by Synthego ICE CRISPR analysis, an indel deconvolution web-based tool (https://synthego.com).

### Cell volume measurement

The Beckman Z2 particle counter and size analyzer was used to determine cell count and volume. The filtering criteria were configured to select cells that were between 10 and 30 μm in size.

### Lysosomal Immunopurification (LysoIP) from mouse tissues and HEK293Ts

Lysosomes were purified from mouse brain and liver as well as HEK293T cells, following established protocols ^17,21^. Mouse brains were obtained post-euthanasia. Small uniform liver samples were collected using a 4 mm diameter biopsy punch to minimize variability between samples. Both the brain and liver tissues were utilized immediately without freezing, employing the steps described in the LysoIP protocols ^17,21^. During the lysosomal lysate enzyme assays, whole brain or liver tissue was sectioned into three pieces and dounced 30X with 3 ml Phosphate Buffer Saline supplemented with protease inhibitor (PBSI). The resulting lysates were centrifuged at 1000*xg* at 4 °C for 10 min. The supernatant was then transferred to a 15 ml falcon tube containing 0.5 ml anti-HA beads and incubated at 4 °C for 1 hour with a rotational mixing. Subsequently, the mixture underwent thorough washing steps and isolation processes to obtain protein fractions suitable for lysosomal lysate experiments as previously described ^10^. Before enzymatic assays were performed, protein concentrations in the lysates were quantified using Pierce BCA Protein Assay kit (Thermo Fisher Scientific). For cell-based assay, an equal number of cells were cultured and allowed to reach at least 80% confluence before each lysoIP. The homogenization and LysoIP from HEK293Ts were adapted from ^17^ and (<u>dx.doi.org/10.17504/protocols.io.bybjpskn</u>).

### Expression and purification of proteins

Expi293F cells were cultured using a combination of 2/3 Freestyle293 and 1/3 Expi293 media and transfected with the PLA2G15 plasmids at a density of 3E6 cells/mL. Using FectoPRO transfection reagent (Polyplus), transfection was performed at a ratio of 1.3 µL FectoPRO to 0.5 µg plasmid DNA per mL of cells. Following transfection, glucose and valproic acid were added right away to reach final concentrations of 0.4% glucose (w/v) and 0.05% valproic acid (w/v). An equivalent amount of valproic acid and glucose were added 1-day post-transfection and cells were harvested 3 days post-transfection. The culture media was harvested and combined with PBS in a 1:1 ratio. HisPur Ni-NTA resin was added to the mixture and incubated at 4 °C for 16 hours. The resin was washed 4 times consecutively using different wash compositions and packed onto a column. The first wash included 50 mM HEPES pH 7.25, 500 mM NaCl, 0.1 mM EDTA, 5 mM beta-mercaptoethanol, 1 mM DTT, 1 mM PMSF, 20 mM imidazole at pH 7.25, cOmplete EDTA-free protease inhibitor cocktail (Roche), 5% glycerol [v/v], and 1% Triton X-100 [v/v]. The second wash was identical to the first wash except for the addition of 10 mM imidazole pH 7.25 and absence of Triton X-100 and protease inhibitor. Imidazole was absent from the third wash, which was identical to the second wash, except that it contained 250 mM NaCl. Except for the addition of 125 mM NaCl, the fourth wash was identical to the third wash. A buffer containing 300 mM imidazole pH 7.25, 50 mM HEPES pH 7.25, 125 mM NaCl, 5% glycerol [v/v], 5 mM BME, 1 mM DTT, 0.1 mM EDTA, 1 mM PMSF and protease inhibitors was used to elute the protein. Using an Amicon 10-kDa MWCO concentrator, the eluted protein was concentrated and further purified by size-exclusion chromatography (SEC) on a Superdex 200 10/300 column. After that, fractions were combined, concentrated using an Amicon 10-kDa MWCO concentrator to achieve a final protein concentration of 1 mg/mL, and swiftly frozen in liquid nitrogen.

### Preparation of liposomes

Unless otherwise specified, 18:1 diether PC (Avanti) was used to make lipid liposomes. The lipids were dried and then combined in a tube with 2 mL water and 3 mL diisopropyl ether to create liposomes that did contain just glycerophospholipids. The resultant mixture was sonicated for 10 min in a water bath to create tiny unilamellar vesicles, which were subsequently dried. Dried liposomes were resuspended in water and Stewart assay was used to determine their concentration.

### Enzyme assays

Recombinant PLA2G15-6xHis at 100 nM concentration was mixed with phospholipid substrates (BMP or PG isomers) of varying concentrations in 50 mM Sodium Acetate:Acetic Acid at pH 4.5 and 150 mM NaCl. The liposomes containing phospholipids had an 18:1 diether PC that was non-cleavable. All lipids and their catalogue numbers are provided in the materials table. 2,2’ *R,R* BMP and 2,2’ *S,R* BMP were obtained from WuXi AppTec. The mixture was incubated for 30 seconds when monitoring products and 10 min when monitoring substrates at 37 °C and then heat-inactivated at 95 °C for 5 min. To adjust for background in each experiment, controls including all components, but the enzyme were also made for each substrate. A time course at various substrate concentrations was carried out to guarantee the linearity of the velocity. The reaction mixture was transferred to plastic autosampler vials to monitor GPG release. In the case of BMP or LPG monitoring, lipids were extracted as described below and transferred to glass autosampler vials. For kinetic studies, corresponding standard for GPG, LPG or BMP was measured in increasing concentrations to extrapolate the concentrations of the products. For time dependent BMP isomer comparison, 100 nM recombinant WT PLA2G15-6xHis was mixed with 20 µM BMP substrates in 50 mM Sodium Acetate:Acetic Acid at pH 4.5 and 150 mM NaCl. The pH optimum was determined by using buffers containing Sodium Acetate:Acetic Acid, HEPES, and Boric acid at their respective pH ranges.

### Enzyme activity assays in lysosomal protein extract

Lysosomes obtained from mouse tissues or HEK293T cells were lysed using hypotonic water for a duration of 1 hour to release soluble lysosomal proteins. Lysosomal protein extract was then combined with 1 µM 3,3’ *S,S* BMP and incubated at 37 °C in 50 mM Sodium Acetate:Acetic Acid pH 5 and 150 mM NaCl for indicated time points. The reaction was then stopped using chloroform:methanol solution in a ratio of 2:1 (v/v) for lipids or 80% methanol for polar metabolite measurements.

### Inhibition assays

For inhibition assays, each inhibitor was preincubated with the substrates before either purified PLA2G15-6xHis or lysosomal lysate was added. The reaction buffer has (50 mM Sodium Acetate:Acetic Acid at pH 5 and 150 mM NaCl) and we followed the enzyme assays described earlier.

### Pulse chase experiments

A 10X concentration of BMP isomers at a concentration of 100 μM in ethanol, along with fatty acid-free BSA at the same concentration in PBS, was initially prepared. This concentrated solution was then diluted to a final concentration of 1X in DMEM (ThermoFisher). *CLN5 PLA2G15* double knockout cells were seeded on each well of a poly-l-lysine coated 6-well plate. Next day, medium was gently aspirated and replaced with serum-free DMEM containing 1X solution of BSA-conjugated BMP and incubated for 2 hours. After incubation, cells were washed once with serum free medium and replaced with complete DMEM containing vehicle solution (1XPBS), 150 nM WT PLA2G15-6XHis or 150 nM PLA2G15(S198A) mutant to chase BMP degradation at varying times. Subsequently, both treated and untreated cells were collected in 200 µL 80% methanol and lipids were extracted using a two-phase extraction method involving a 2:1 chloroform:methanol (v/v) mixture as described below. Finally, lipids were analyzed with LC-MS.

### Enzyme supplementation

To validate the lysosomal localization of recombinant enzymes, PLA2G15-6xHis and mutant were labeled with NHS-Alexa488 dye (Thermo Fisher Scientific) according to the manufacturer’s protocol. Afterwards, 50,000 of *PLA2G15* knockout or *CLN5 PLA2G15* double knockout cells were seeded with 700 µl DMEM medium on each division of a 4-chamber 35 mm glass bottom dish (Cellvis). Then, cells were treated with 50 nM fluorescently labeled protein in complete media for 2 days. After 2-day incubation, medium was replaced with 700 µl of DMEM medium containing 1 µM LysoTracker Red DND-99 (ThermoFisher) and 1 µg/ml Hoechst 33342 and incubated for 30 min. Medium were then replaced again with fresh medium and immediately imaged with Leica LSM980 confocal microscope and images were analyzed using ImageJ.

For enzyme supplementation experiments, 600,000 cells were seeded on a 6-well plate. After 24 hours, cells were treated with 50 nM PLA2G15 wildtype and mutant proteins in complete media. After 48 hours, lipids were extracted as described above and analyzed by lipidomics.

### Lipid and polar metabolite extraction

Samples were processed for lipid or polar metabolite extraction following previous protocol ^21^. For lipid extraction, 950 µL of 2:1 (v/v) chloroform:methanol containing SPLASH LipidoMIX internal standard mix at a concentration of 750 ng/ml was added to all samples. The mixture was vortexed for 1 hour at 4 °C. Subsequently, 200 µL of 0.9% (w/v) NaCl was added and vortexed for 10 min at 4 °C. Then, the mixture was spun at 3,000*xg* for 15 min at 4 °C. The lower chloroform phase, which contained the lipids, was collected and dried using a SpeedVac. Next, the dried lipid extracts were reconstituted in 50 µL of 13:6:1 (v/v/v) acetonitrile(ACN):isopropyl alcohol:water and vortexed for 10 min at 4 °C in a cold room. Then, the samples were centrifuged at max speed for 10 min at 4 °C and 45 µL transferred into glass vials with insert for LC-MS analysis. For GPG analysis, samples were extracted using 80% methanol in LC-MS grade water containing 500 nM isotope-labeled amino acids as internal standards (Cambridge Isotope Laboratories). Samples were then vortexed for 10 min at 4 °C, centrifuged at 20,627*xg* and transferred into polar vials for LC-MS.

### Untargeted lipidomics analysis

Lipidomic analysis of samples in Extended Data Fig. 9h,i and Supplemental Table 8 was performed at the Core Facility Metabolomics of the Amsterdam UMC, Amsterdam, the Netherlands. In a 2 mL tube, the following amounts of internal standards dissolved in 1:1 (v/v) methanol:chloroform were added to each sample: Bis(monoacylglycero)phosphate BMP(14:0/14:0) (0.2 nmol), Ceramide-1-phosphate C1P (d18:1/12:0) (0.125 nmol), D7-Cholesteryl Ester CE(16:0) (2.5 nmol), Ceramide Cer(d18:1/12:0) (0.125 nmol), Ceramide Cer(d18:1/25:0) (0.125 nmol), Cardiolipin CL(14:0/14:0/14:0/14:0) (0.1 nmol), Diacylglycerol DG(14:0/14:0) (0.5 nmol), Glucosylceramide GlcCer(d18:1/12:0) (0.125 nmol), Lactosylceramide LacCer(d18:1/12:0) (0.125 nmol), Lysophosphatidic acid LPA(14:0) (0.1 nmol), Lysophosphatidylcholine LPC(14:0) (0.5 nmol), Lysophosphatidylethanolamine LPE(14:0) (0.1 nmol), Lysophosphatidylglycerol LPG(14:0) (0.02 nmol), Phosphatidic acid PA(14:0/14:0) (0.5 nmol), Phosphatidylcholine PC(14:0/14:0) (2 nmol), Phosphatidylethanolamine PE(14:0/14:0) (0.5 nmol), Phosphatidylglycerol PG(14:0/14:0) (0.1 nmol), Phosphatidylinositol PI(8:0/8:0) (0.5 nmol), Phosphatidylserine PS(14:0/14:0) (5 nmol), Sphinganine 1-phosphate S1P(d17:0) (0.125 nmol), Sphinganine-1-phosphate S1P(d17:1) (0.125 nmol), Ceramide phosphocholines SM(d18:1/12:0) (2.125 nmol), Sphingosine SPH(d17:0) (0.125 nmol), Sphingosine SPH(d17:1) (0.125 nmol) and Triacylglycerol TAG(14:0)_2_ (0.5 nmol). All internal standards were purchased from Avanti Polar Lipids, Alabaster, AL. After the addition of the internal standards, 1.5 mL 1:1 (v/v) methanol:chloroform was added before thorough mixing. The samples were then centrifuged for 10 min at 14.000 rpm, supernatant transferred to a glass vial and evaporated under a stream of nitrogen at 60°C. The residue was dissolved in 150 μL of 1:1 (v/v) methanol:chloroform. Lipids were analyzed using a Thermo Scientific Ultimate 3000 binary HPLC coupled to a Q Exactive Plus Orbitrap mass spectrometer. For normal phase separation, 5 μL of each sample was injected onto a Phenomenex® LUNA silica, 250 * 2 mm, 5µm 100Å. Column temperature was held at 25°C. Mobile phase consisted of (A) 85:15 (v/v) methanol:water containing 0.0125% formic acid and 3.35 mmol/l ammonia and (B) 97:3 (v/v) chloroform:methanol containing 0.0125% formic acid. Using a flow rate of 0.3 ml/min, the LC gradient consisted of: isocratic at 10% A 0-1 min, ramp to 20% A at 4 min, ramp to 85% A at 12 min, ramp to 100% A at 12.1 min, isocratic at 100% A 12.1-14 min, ramp to 10% A at 14.1 min, isocratic at 10% A for 14.1-15 min. For reversed phase separation, 5 μL of each sample was injected onto a Waters HSS T3 column (150 x 2.1 mm, 1.8 μm particle size). Column temperature was held at 60°C. Mobile phase consisted of A 4:6 (v/v) methanol:water and B 1:9 (v/v) methanol:isopropanol, both containing 0.1% formic acid and 10 mmol/L ammonia. Using a flow rate of 0.4 mL/min, the LC gradient consisted of: isocratic at 100% A at 0 min, ramp to 80% A at 1 min, ramp to 0% A at 16 min, isocratic at 0% A for 16-20 min, ramp to 100% A at 20.1 min, isocratic at 100% A for 20.1-21 min. MS data were acquired using negative and positive ionization by continuous scanning over the range of m/z 150 to m/z 2000 at a resolution of 280,000 full width at half maximum. Data were analyzed using an in-house developed lipidomics pipeline written in the R programming language (http://ww.r-project.org) and MATLAB. Lipid identification was based on a combination of accurate mass, (relative) retention times, fragmentation spectra (when required), analysis of samples with known metabolic defects, and the injection of relevant standards. Lipid classes are defined in our lipidomics pipeline in terms of their generic chemical formula, where R represents the radyl group. Upon import of the lipid database in the annotation pipeline the generic chemical formula of each lipid class is expanded by replacing the R-element with a range of possible radyl group lengths and double bond numbers. The resulting expanded list of chemical formulas is then used to calculate the neutral monoisotopic mass of each species. The reported lipid abundances are semi-quantitative and calculated by dividing the response of the analyte (area of the peak) by that of the corresponding internal standard multiplied by the concentration of that internal standard (arbitrary unit, A.U). The suffix [O] indicates lipids containing an alkyl-ether group, whereas the addition of a quote ‘ indicates an alkenyl-ether group. As no dedicated internal standard for ether lipids are available, we used the (L)PC/PE internal standards to normalize the corresponding ether lipid species as described above.

### Targeted lipidomics and polar metabolomics

Targeted analyses were adapted and performed as previously described ^2,10,39^. Lipids were separated on an Agilent RRHD Eclipse Plus C18,2.1×100mm,1.8u-BC column with an Agilent guard holder (UHPLC Grd, Ecl. Plus C18,2.1mm,1.8µm) while polar metabolites were separated using hydrophilic interaction chromatography (HILIC) using a SeQuant® ZIC®-pHILIC 50 x 2.1 mm column (Millipore Sigma 1504590001) with a 20 x 2.1 mm (Millipore Sigma 1504380001) guard. Before mass spectrometry, the columns were connected to a 1290 LC system for lipid and metabolite separation. The LC system was linked to an Ultivo triple quadrupole (QQQ) mass spectrometer with an LC-ESI (electrospray ionization) probe. External mass calibration was done weekly using a QQQ standard tuning mix. The column compressor and autosampler were held at 45 °C and 4 °C, respectively. For lipidomics, the mass spectrometer parameters included a capillary voltage of 4.4 kV in positive mode and 5.5 kV in negative mode, and gas temperature and sheath gas flow were held at 200 °C and 275 °C, respectively. Both gas flow and sheath gas flow were 10 L/min and 11 L/min, respectively, while the nebulizer was maintained at 45 psi. The nozzle voltages were maintained at 500 in positive mode and 1000 in negative mode. These conditions were held constant for both ionization modes acquisition. For metabolomics, similar conditions were used except gas temperature and sheath gas flow were held at 250 °C and 300 °C, respectively. Additionally, gas flow and nebulizer were 13 L/min and 30 psi.

For lipid measurements, 2-4 µL injection volumes were used for each sample for polarity switching. A mobile phase with two distinct components, A and B, was employed for the chromatographic process. Mobile phase A was a mixture of acetonitrile: water (2:3 [v/v]), while mobile phase B was composed of isopropanol: acetonitrile (9:1 [v/v]), both also containing 0.1% formic acid and 10 mM ammonium formate. The elution gradient was carried out over a total of 16 min, with an isocratic elution of 15% B for the first min, followed by a gradual increase to 70% of B over 3 min, and then to 100% of B from 3 to 14 min. Subsequently, this was maintained from 14 to 15 min, after which solvent B was decreased to 15% and maintained for 1 min plus 2 min to allow for column re-equilibration. The flow rate was set at 0.400 mL/min.

For polar metabolite measurements, each sample was injected with a volume of 1-2.5 µl. Mobile phase A is made of 20 mM ammonium carbonate and 0.1% ammonium hydroxide dissolved in LC/MS grade water, while B is composed of 100% LC/MS grade acetonitrile. Elution was performed over a 10-min gradient; the B component was linearly decreased from 80% to 20% between 0-7 min, rapidly increased from 20-80% between 7-7.5 min, and was held at 80% B from 7.5-10 min. The flow rate was set to 0.15 mL/min.

The QQQ was set to operate in multiple reaction monitoring (MRM) to analyze compounds of interest. Standard lipids, glycerophosphodiesters (GPDs) and amino acids were optimized using MassHunter Optimizer MRM, a software used for automated method development. For most species, the two most abundant transitions were selected to detect it. The precursor-product ion pairs (m/z) of the compounds used for MRM are listed in Supplementary Table 10.

Polar metabolites and lipids were annotated, and their abundances were measured using the Qualitative MassHunter acquisition software and QQQ quantitative analysis software (Quant-My-Way), respectively. The peak areas of all metabolites were integrated using the retention time and MRM method, and the resulting raw abundances were exported to Microsoft Excel for further analysis. The raw abundances of the internal controls, endogenous amino acids and highly abundant lipids were examined to ensure accuracy of the analysis. Raw abundances were normalized to cell number using abundance of endogenous control lipids in the same sample. Quality control samples were consistently measured to ensure optimal instrument performance and linearity. The average of the blank samples was subtracted from all raw abundances. For fold change calculations, each sample was divided by the average of the wildtype or control samples.

### Cholesterol measurement

Cholesterol was detected in left hemibrain and left liver lobe from 4 animals per group (3 samples each from each animal) using a commercial kit (ab65359; Abcam). Left hemibrains as well as left liver lobe were homogenized in 4 v/w PBS using UPHO beadmill (Geneye, SOP NEQU200) at 60Hz for 55sec. 100 µl of the homogenate were transferred to a fresh Eppendorf tube for further processing, one 5 µl aliquot was used for total protein determination (Pierce™ BCA assay, Thermo Fisher Scientific, 23225). For extraction 200 µl of Chloroform: Isopropanol: NP-40 (7:11:0.1) were added to the 100 µl aliquot and the sample was vortexed vigorously two times after 5-10 min incubation phase in between. Thereafter, the extract was spun 5 – 10 minutes at 15,000 x g in a centrifuge. The entire chloroform phase was transferred to a new tube and air dried at 50 °C up to 24h. The dried lipids were dissolved by vortexing with 100 μl of Assay Buffer II (from kit ab65359) and stored at −80 °C until further use. Cholesterol was assessed in parallel in all samples following the protocol in the commercially available Cholesterol/ Cholesteryl Ester Quantitation Assay Kit (ab65359). Fluorescence was measured on a microplate reader (Cytation 5, SOP NEQU201) at Ex/Em = 535/587 nm.

### Immunoblotting

Purified proteins were run on SDS-PAGE (Thermo Fisher Scientific) at 120V and transferred to PVDF membranes at 40V for 2 hours. The membranes were then blocked with 5% non-fat dry milk in Tris-buffered saline with 0.1% Tween 20 (TBST) for 30 min and incubated overnight with primary antibodies in 5% BSA in TBST at 4 °C using dilutions listed in the antibody table. After incubation, membranes were washed 3X with TBST for 5 min per wash and then incubated with appropriate secondary antibodies diluted 1:3,000 in 5% BSA for 1 hour at room temperature. Membranes were then washed 3X with TBST followed by ECL2 western blotting substrate (Thermo Fisher Scientific) prior to visualization.

### Thermal Stability Assay

A Tycho NT.6 instrument (Nanotemper Technologies) was used to determine the melting stability of human PLA2G15-6xHis wildtype and S198A. A concentration of 5 µM purified proteins in 50 mM Sodium Acetate-Acetic Acid (pH=5.0) and 150 mM NaCl were loaded into Tycho NT.6 capillaries. The melting temperature (Tm) was obtained by plotting the absorbance ratio of solvent-exposed tryptophan vs buried tryptophan (350 nm/330 nm) throughout a temperature gradient and determining the inflection temperature from the first derivative.

### Microscale Thermophoresis (MST)

Recombinant human PLA2G15-6xHis S198A were labeled using RED-NHS Protein Label Kit following the manufacturer’s protocol. Monolith NT.115 instrument was used for binding experiments. A 100 nM labeled protein was incubated with serial dilutions of substrates in 50 mM Sodium Acetate-Acetic Acid (pH=5.0), 150 mM NaCl, and 5 µM BSA for 30 min at room temperature. Using premium capillaries, binding measurements were conducted at 20%, 40%, or 60% MST power for 30 s with 5 s of cooling. The dissociation constant Kd was derived by plotting the fraction bound against logarithmic substrate concentrations according to the law of mass action.

### Synthesis of GPG

Standard GPG was prepared for the kinetic assays. Phosphatidylglycerol (18:1/18:1 PG) (Avanti) was saponified to produce GPG according to the following protocol: 100 mg of PG was dissolved in a 20 ml scintillation vial that contained a 12 ml solution of 2:1 chloroform:methanol with a magnetic stirring bar. NaOH (4 ml of 2 M) was added to the mixture and the reaction was stirred for 2 hours at room temperature. To neutralize the reaction, HCl (4 ml of 2 M) was used, and GPG was extracted with 8 ml water into separate vessels twice. After mixing the aqueous phases, they were frozen in liquid nitrogen, lyophilized, and cleaned with 8 mL of chloroform. To separate the crude product from inorganic salts, lyophilates were dissolved in a minimal amount of methanol. The desired products were then extracted from the precipitate using paper filtering.

### Thin Layer Chromatography (TLC)

100 nM of PLA2G15-6xHis WT was mixed with 20 µM BMP and incubated at 37 °C for 1 hour as described for the enzyme assay. Two microliters of reaction were spotted on silica and optimized to observe product release. The silica was placed in a thin layer chromatography (TLC) solvent (CHCl_3_/CH_3_OH/conc ammonia/water (90:54:5.5:2 [v/v]) and dipped into an iodine stain (Iodine crystals and silica gel mixture) after the solvent run prior to visualization.

### Diffdock Docking

Structure of *H. Sapiens* PLA2G15 was obtained from the AlphaFold entry of Q8NCC3 (UniProt). The predicted protein structure was prepared for docking using ChimeraX to add hydrogens according to the expected protonation state at pH 5. The substrates were drawn in ChemDraw to extract the SMILES string. The structure of the protein (pdb format) and the SMILES string were used as input for DiffDock. We ran DiffDock in a web server using the following parameters: twenty inference steps, eighteen actual inference steps, and forty samples. The docking results with the best confidence score were used for the analysis using ChimeraX.

### siRNA knockdown in mouse BMDMs

100,000 of BMDMs isolated from *Gba^+/+^* or *Gba*^+/D409V^ mice were seeded on each well on each division of a 4-chamber 35 mm glass bottom dish (Cellvis) that contains 750 µl complete DMEM. Next day, cells were transfected with scramble siRNA (ThermoFisher 4390844) or two siRNA targeting mouse Pla2g15 (ThermoFisher s101347, s101348) with RNAiMax (ThermoFisher) according to the manufacturer’s instruction. 3 days post transfection, cells were subjected to LysoFQ assay as described below.

### LysoFQ-GCase activity assay

About 50,000 of bone marrow derived macrophages were seeded with 700 µL macrophage medium on each division of a 4-chamber 35 mm glass bottom dish (Cellvis). In non-siRNA experiment, 300 nM recombinant PLA2G15 wildtype or mutant or vehicle solution (1X PBS) was added during cell seeding. After overnight incubation, 10 µM of LysoFQ-GBA probe ^69^ was added to the medium and incubated for 30 min. After incubation, medium was washed once with macrophage medium and then replaced with fresh medium containing 100 nM AT3375 (GCase inhibitor) to stop the reaction. Cells were then imaged with Leica LSM confocal microscope and images were analyzed using image J.

### Haploid genetic screens

To identify regulators of intracellular cholesterol levels we prepared libraries of mutagenized Hap1 wild-type and NPC1-deficient cells using a gene-trap retrovirus expressing blue fluorescent protein (BFP), as described previously ^70^. For each genotype, mutagenized cells were cultured and expanded for one week after mutagenesis. Subsequently, cells were washed with PBS, dissociated, pelleted, and fixed with BD Fix buffer I (BD Biosciences) for 10 min at 37 °C. After washing once with FACS buffer (PBS containing 1% BSA) and once with PBS, the cells were pelleted and resuspended in permeabilization buffer (PBS containing 0.1% saponin). Cells were incubated for 5 min at room temperature while rotating. Cells were pelleted, washed twice with FACS buffer, and incubated with FACS buffer containing 3 µg/ml AlexaFluor647-labeled Perfringolysin O (PFO) ^71^ and 1 µg/ml 4′,6-diamidino-2-phenylindole (DAPI) for 45 min at room temperature while rotating. Cells were subsequently pelleted, washed twice with FACS buffer before sorting two populations of cells (i.e. PFO-LOW and PFO-HIGH) that represent ∼5% of the lowest and highest cholesterol containing cells from the total cell population, respectively. In addition, cells were sorted in parallel for haploid DNA content (G1 phase) using the DAPI signal. Cell sorting was carried out on a FACS Aria III cell sorter (BD Biosciences) until ∼10 million cells of each population were collected. Sorted cells were pelleted and genomic DNA was isolated. Gene-trap insertion sites of each sorted cell population were amplified and mapped by deep sequencing. For each gene in a single screen, a mutation index (MI) was calculated corresponding to the ratio of the number of disruptive integrations per gene in both populations normalized by the number of total integrations in each channel (i.e. PFO-LOW and PFO-HIGH) as previously described ^70^. Genes significantly enriched for disruptive gene-trap integrations in either the PFO-HIGH or PFO-LOW query populations were identified using a two-sided Fisher’s exact test. Resulting P values were adjusted for multiple testing using the Benjamini–Hochberg false discovery rate correction.

### siRNA knockdown in NPC1 patient fibroblast

Two NPC1 patient fibroblast cells (GM03123 and GM18453) were acquired from Coriell with agreement. 30,000 cells of two NPC1 patient fibroblasts were seeded on each well of an 8-chamber cell culture slide (Celltreat, Pepperell, MA) that contains 500 µL complete DMEM. Next day, cells were transfected with scrambled siRNA (ThermoFisher 4390844) or two siRNAs targeting human PLA2G15 (ThermoFisher s24296, s24297) using RNAiMax (ThermoFisher) according to the manufacturer’s instruction. 2 days post transfection, medium was gently aspirated, and cells were washed once with 800 µL PBS, followed by immediate 15 min fixation with 200 µL of 4% PFA in PBS. Then, cells were washed once with 800 µL PBS and stained with 200 µl of 1 mg/ml Filipin (Cayman). Slides were sealed with glass coverslip and mounting solution (Thermo Scientific) and imaged with Leica LSM980 confocal microscope.

### Data preparation and statistics

For quantitative data presentation, GraphPad Prism v.10.2 was used and figures were assembled in Adobe Illustrator 2023. All statistical analyses were performed in Prism. Unless otherwise stated in the legends, all measurements presented were derived from independent samples or biological replicates. Western blot data displayed were representative experiments. BioRender was used with permission to illustrate the figure schematics for Fig. 1g, 4b and Extended Data Fig. 6d. ChemDraw 22.2.0 was used in chemical structures depicted in Fig. 1a-c, 2a and Extended Data Fig. 5h. Extended Data Fig. 9e was adapted from images provided by Servier Medical Art (Servier; https://smart.servier.com/), licensed under a Creative Commons Attribution 4.0 Unported License.

## ACKNOWLEDGEMENT

We thank all members of the Abu-Remaileh laboratory and of Scenic Biotech for helpful discussions. We also thank Suzanne Pfeffer for providing MST and James A. Shayman for PLA2G15-deficient mouse model. Fred Vaz from UMC Amsterdam for providing untargeted lipid analysis and services. The *Npc1^m1N/J^* mouse model work was performed at Scantox, Vienna, Austria.

## Author Contributions

Conceptualization: K.N and M.A-R. Methodology: K.N., J.X., and M.A-R. Investigation: K.N., J.X., H.N.A., and I.V.P. Mass spectrometry data acquisition and analysis: K.N. J.A.S. provided PLA2G15-deficient tissues. Histopathology: R.d.M. Genetic modifier screen: M.R., V.A.B., S.M.B.N., and T.R.B. Pla2g15 and Pla2g15/Npc1 mice: M.R., A.d.J., V.A.B., S.M.B.N., T.R.B., and G.H. coordinated activities. Supervision: M.A-R. Manuscript writing-original draft: K.N. and M.A-R. All authors reviewed the manuscript. Funding acquisition: M.A-R.

## Data and materials availability

All data needed to evaluate the conclusions stated in the paper are present in the paper and/or the Supplementary Materials. This paper does not report original code. Cell lines can be requested, and requests will be fulfilled by, the lead author, Monther Abu-Remaileh. All sequencing datasets from the haploid screens have been deposited in the NCBI Sequence Read Archive under accession number PRJNA1177366. Data from untargeted lipid profiling are deposited at Zenodo under accession number 14170856.

## Supplementary Information

### Extended Data Figures

**Extended Data Fig. 1:**
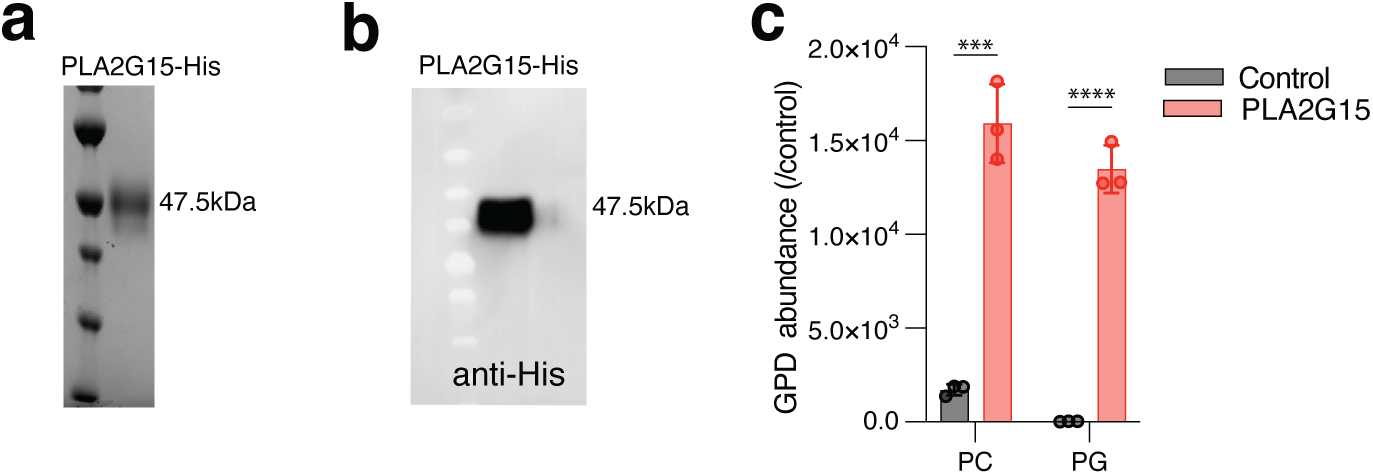
Purification of recombinant PLA2G15. **(a)** Purification of wildtype recombinant His-tagged lysosomal phospholipase PLA2G15. Representative image of 47.5 kDa PLA2G15-6xHis protein purified to homogeneity and stained with Coomassie Blue. **(b)** Immunoblot analysis of purified PLA2G15 using anti-His antibody. **(c)** Purified PLA2G15 is active enzyme. Phospholipase activity determined using 100 nM PLA2G15 against phosphatidylcholine (PC) or phosphatidylglycerol (PG) under acidic conditions for 1 minute. Glycerophosphodiester products (GPD) detected using LC-MS. *n* = 3 independent replicates for each protein. Statistical analysis was performed using two-tailed unpaired *t*-tests. ***p < 0.001 and ****p < 0.0001. Control has all reaction components with no enzyme.

**Extended Data Fig. 2:**
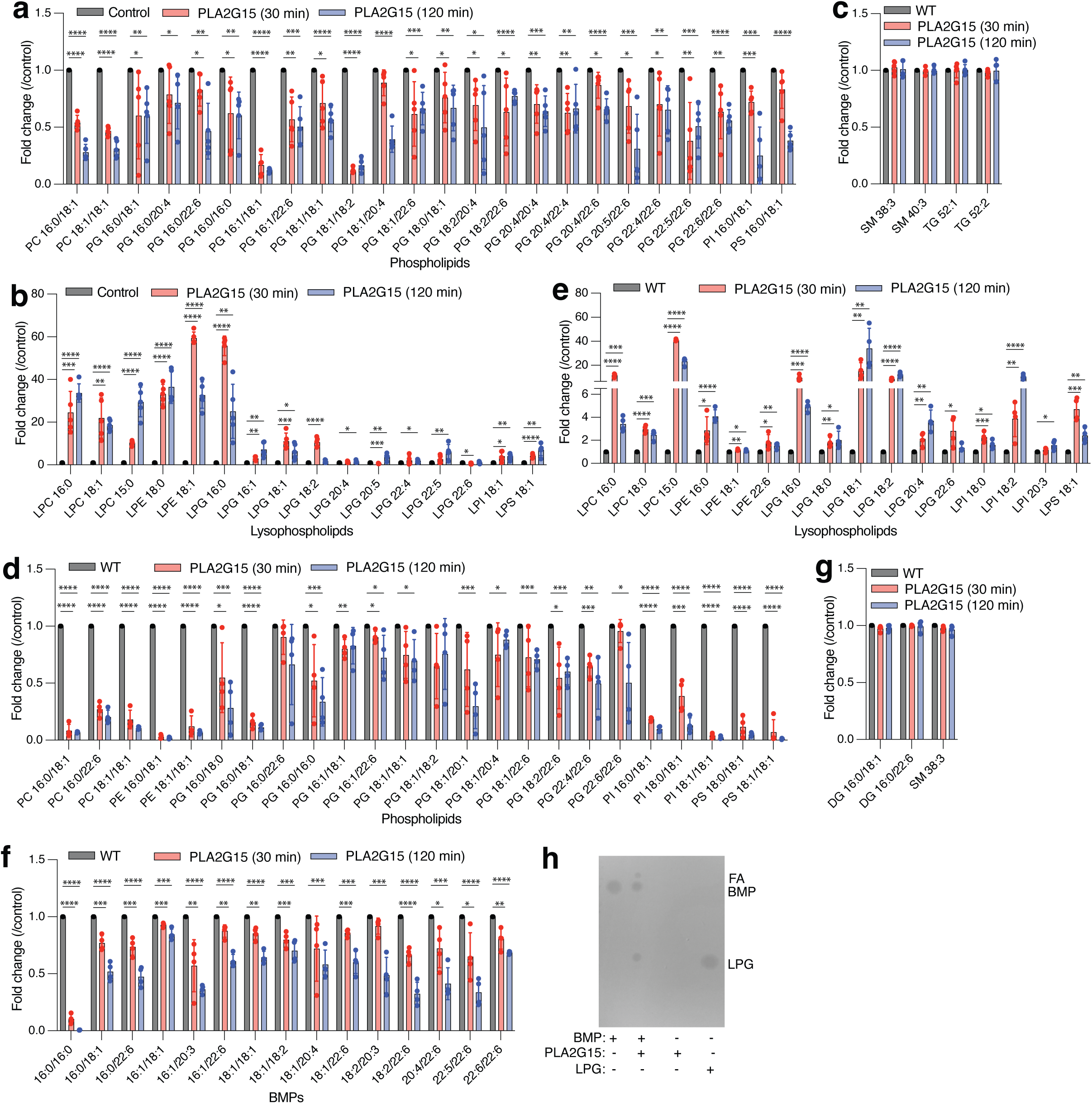
PLA2G15 catabolizes lysosomal phospholipids including BMP. **(a and b)** PLA2G15 hydrolyzes phospholipid substrates isolated from mouse liver lysosomes. Fold changes in the abundance of known PLA2G15 substrates (PC, PE, PG, PI and PS) in (a) and lysophospholipid intermediates (LPC, LPE, LPG, LPI, LPS) in (b) for 30 min or 120 min reactions **(c)** Non-phospholipids are unchanged in PLA2G15 reaction. Fold changes in sphingomyelin (SM) and triacylglycerol (TG). Each incubation time was compared to control (no enzyme) samples from the same mouse incubated for the same time. Data are mean ± SD of *n* = 5 biological replicates. Statistical analysis was performed using two-tailed unpaired *t*-tests. *p < 0.05, **p < 0.01, ***p < 0.001 and ****p < 0.0001. **(d-f)** PLA2G15 hydrolyzes BMP and other phospholipid substrates extracted from HEK293T lysosomes. Lysosomal lipids were isolated from HEK293T lysosomes and incubated with recombinant PLA2G15 for 30 min or 120 min. Fold changes in phospholipid substrates (PC, PE, PG, PI and PS) in (d), lysophospholipid intermediates (LPC, LPE, LPG, LPI, LPS) in (e), and BMPs in (f). **(g)** Non PLA2G15 substrates remained unchanged. Fold changes in diacylglycerol (DG) and sphingomyelin (SM). Each incubation time was compared to control samples from each biological replicate incubated at the same time. Data are mean ± SD of *n* = 4 biological replicates. Statistical analysis was performed using two-tailed unpaired *t*-tests. *p < 0.05, **p < 0.01, ***p < 0.001 and ****p < 0.0001. **(h)** PLA2G15 hydrolyzes synthesized BMP in vitro. Thin Layer Chromatography (see methods) was used to measure the presence of LPG intermediate after incubating wildtype recombinant PLA2G15 with 3,3’ 18:1-*S,S* BMP under acidic conditions (pH=5.0). Controls had similar reaction buffers except with no enzyme. Putative FA: fatty acid was annotated based on solvent characteristics.

**Extended Data Fig. 3:**
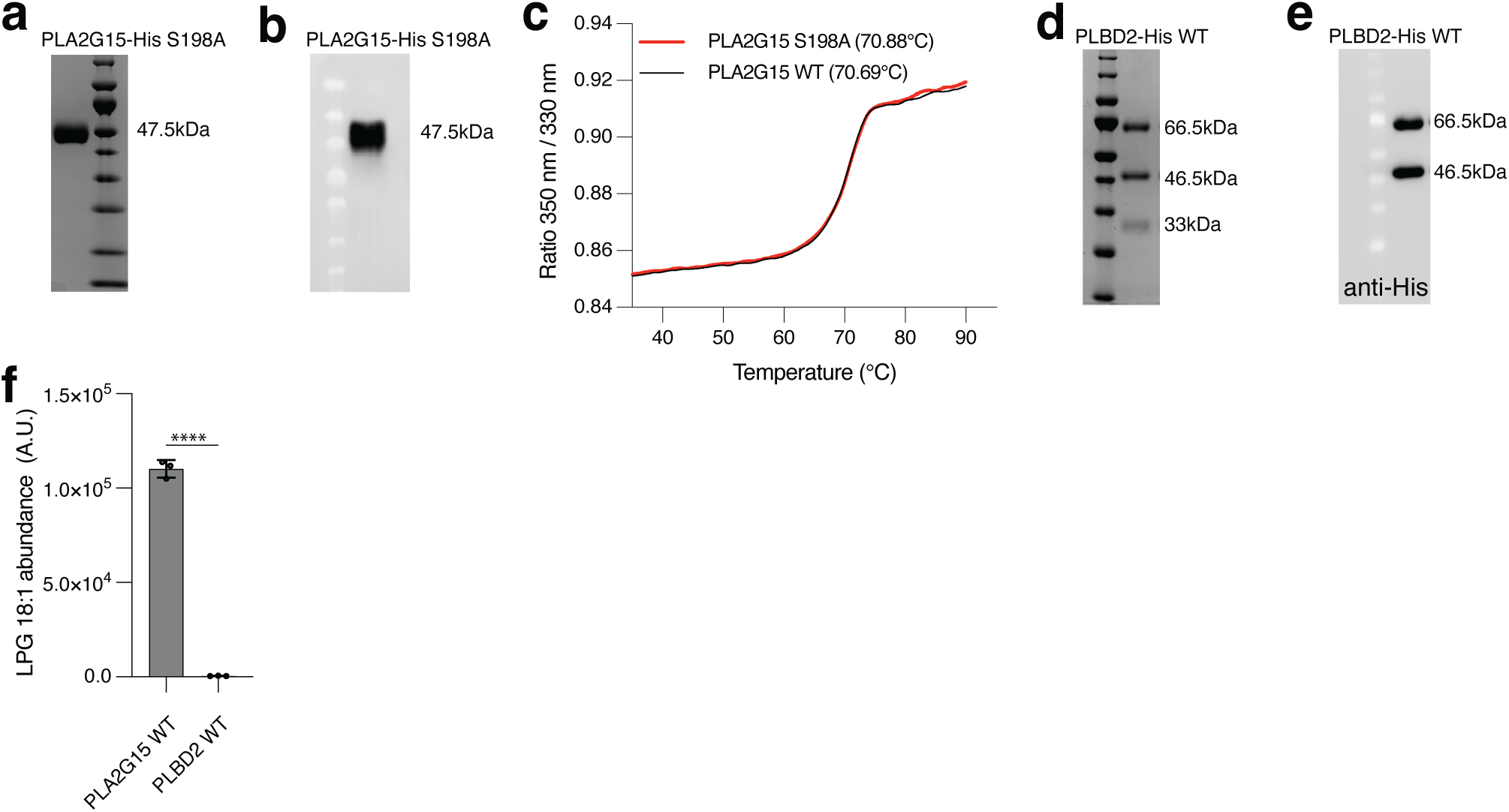
Purification and characterization of lysosomal phospholipases. **(a)** Purification of recombinant His-tagged PLA2G15 S198A. Representative image of 47.5 kDa PLA2G15 S198A-6xHis protein purified to homogeneity and stained with Coomassie Blue. **(b)** Immunoblot analysis of PLA2G15 S198A using anti-His antibody. (c) Melting temperature (Tm) curves indicate that wildtype and S198A mutant maintain thermal stability. Tm is indicated on the graph. **(d)** Purification of recombinant His-tagged PLBD2. Representative image of PLBD2-6xHis protein purified to homogeneity and stained with Coomassie Blue. The three bands are indicative of 66.5 kDa proenzyme, 46.5 kDa matured form and 33 kDa N-terminus pro-domain. **(e)** Immunoblot analysis of PLBD2 using anti-His antibody. **(f)** PLBD2 has no BMP hydrolase activity. BMP Hydrolase activity by 100 nM wildtype recombinant PLA2G15 and PLBD2 proteins under acidic conditions determined using LC-MS. N = 3 independent replicates from each protein.

**Extended Data Fig. 4:**
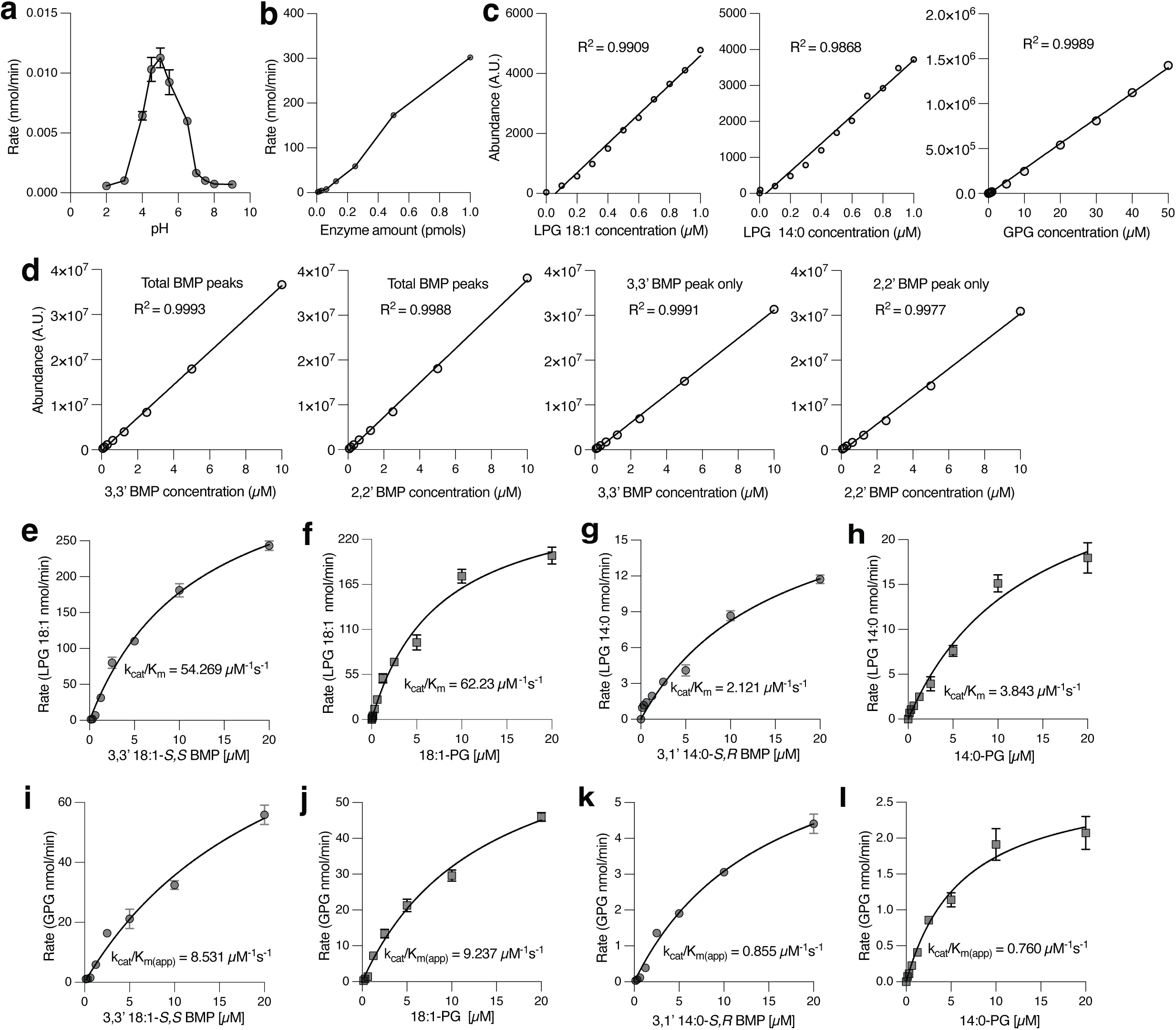
PLA2G15 has high catalytic activity towards BMP under optimized enzyme reaction conditions. **(a)** BMP hydrolase activity of PLA2G15 has an acidic pH optimum. PLA2G15-6xHis was incubated with 3,3’ 18:1-*S,S* BMP under the indicated pH conditions. A representative graph is shown for an experiment repeated at least 3 times. **(b)** PLA2G15-6xHis was incubated with 1 µM of 3,3’ 18:1-*S,S* BMP and at different time points and enzyme concentrations. A representative graph indicates linearity in increments across multiple enzyme concentrations at a 30 second time point. **(c)** Standard curves generated from measuring the indicated LPG and GPG standards. **(d)** Standard curves generated from measuring the indicated BMP standards. **(e-h)** PLA2G15 deacylates BMP to LPG with high catalytic efficiency that is comparable to that against PG phospholipid and this activity correlates with acyl chain length. Recombinant PLA2G15 was incubated with the indicated phospholipids under acidic conditions (pH= 5.0) for 30 seconds and reactions were stopped by heating. Phospholipids were incorporated in liposomes with non-cleavable 18:1 diether PC and indicated LPG intermediates were measured. Each experiment was repeated at least three times, and a representative graph is shown. K_cat_/K_m_ is a measure of kinetic efficiency. **(i-l)** PLA2G15 deacylates BMP to GPG with high efficiency. Recombinant PLA2G15 was incubated with the indicated phospholipids under acidic conditions (pH= 5.0) for 30 seconds and reactions were stopped by heating. Phospholipids were incorporated in liposomes with non-cleavable 18:1 diether PC and GPG product was measured. Each experiment was repeated at least three times, and a representative graph is shown. “app” is defined as apparent because the catalytic efficiency is derived from a two-step reaction.

**Extended Data Fig. 5:**
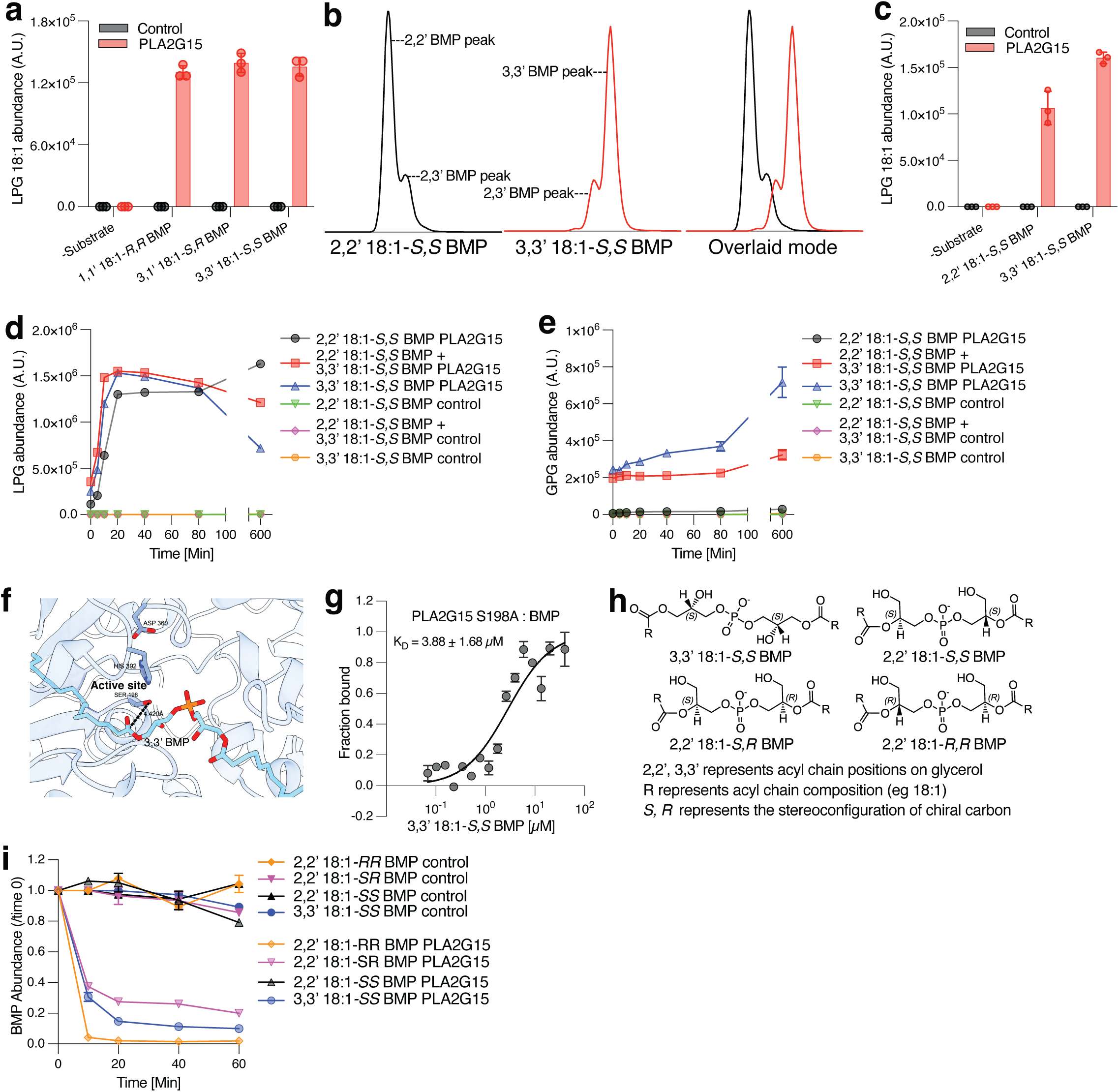
*sn2, sn2’* esterification position of *S,S* BMP confers resistance to PLA2G15-mediated hydrolysis in vitro. **(a)** PLA2G15 efficiently deacylates BMP independent of stereoconfiguration. Hydrolysis of 1 µM BMP stereoisomers by 100 nM recombinant PLA2G15 for 30 seconds under acidic conditions (pH=5.0). LPG intermediate was measured with LC-MS. Data are mean ± SD of *n* = 3 independent replicates. **(b)** A representative image of the extracted ion chromatogram of 2,2’ and 3,3’ positional BMP isomers used in this paper. Each compound has a minor amount of 2,3’ isomer. **(c)** LPG monitoring shows minimal difference in the activity of PLA2G15 against positional BMP isomers. Same experiment as in (a) using 1 µM of either 3,3’ *S,S* BMP or 2,2’ *S,S* BMP as substrate incubated for 30 secs. Data are mean ± SD of *n* = 3 independent replicates. **(d and e)** Time course of BMP hydrolysis shows 2,2’ BMP positional isomer confers resistance to PLA2G15 degradation. Hydrolysis of 20 µM 2,2’ *S,S* BMP or 3,3’ *S,S* BMP positional isomers as well as equimolar amount of both by 100 nM recombinant PLA2G15 over 10 hours under acidic conditions (pH=5.0). Controls had similar reaction buffers except with no enzyme. BMP hydrolase activity was determined using LC-MS by measuring the abundance of LPG intermediate in (d) and GPG product in (e). Data are mean ± SD of *n* = 3 independent replicates. **(f)** DiffDock of 3,3’ *S,S* BMP onto AlphaFold human PLA2G15 structure. Catalytic triad residues Ser198, Asp360 and His392 of the active site of PLA2G15 are annotated. **(g)** BMP binds to PLA2G15 S198A at low micromolar concentration. Binding affinity was determined using microscale thermophoresis for 3,3’ *S,S* BMP. Data presented as mean ± SEM from three measurements. **(h)** Chemical structures of other BMP lipids used in this study. **(i)** Time dependent hydrolysis of BMP stereoisomers confirms *S,S* stereochemistry of 2,2’ BMP contributes to resistance to PLA2G15-dpendent degradation. Assay as in (d) using indicated BMP isomers over 1 hour under acidic conditions (pH=5.0). Data are mean ± SD of *n* = 3 independent replicates.

**Extended Data Fig. 6:**
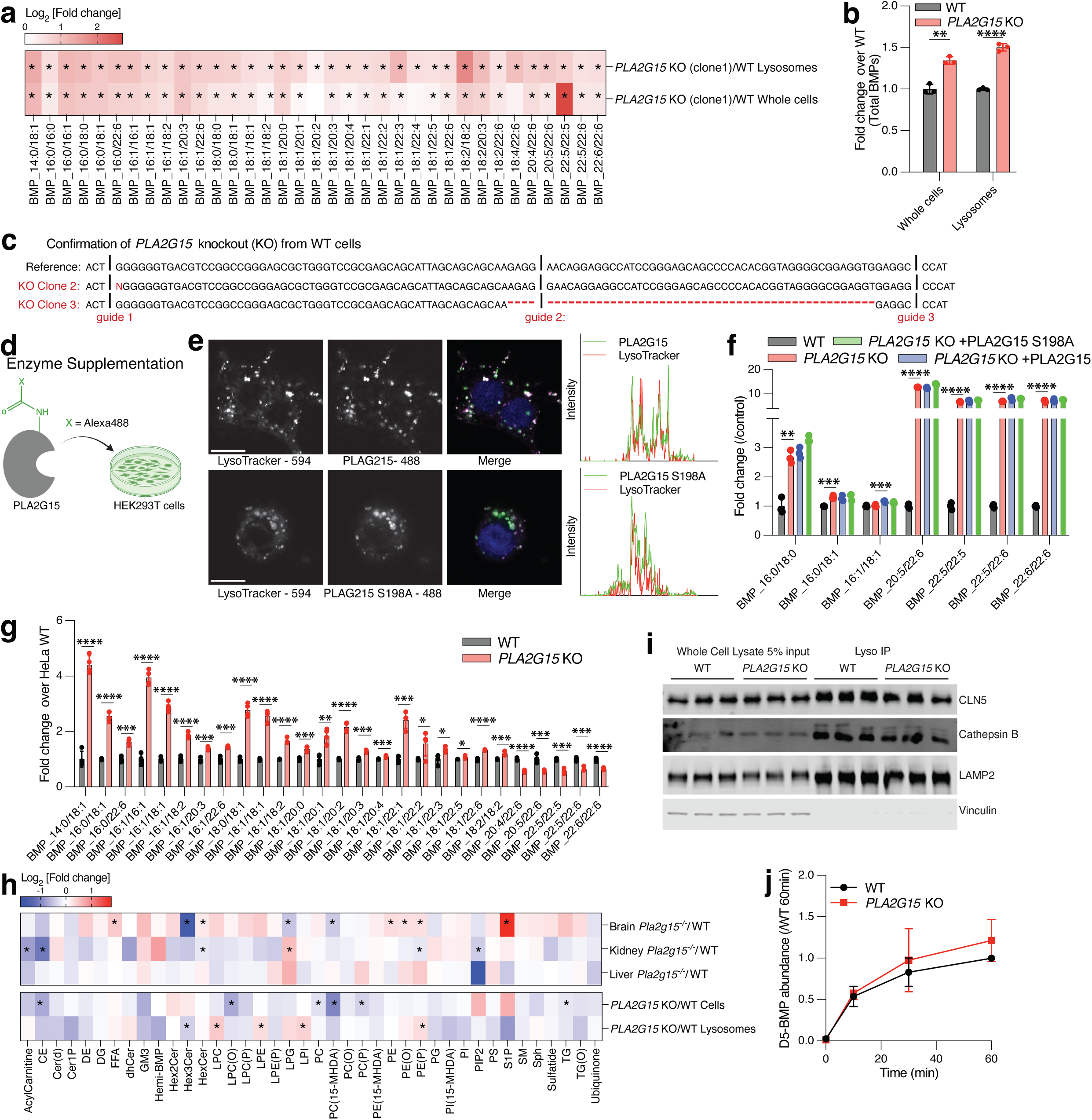
PLA2G15-deficient cells and lysosomes accumulate BMP. **(a and b)** Targeted analyses of BMP lipids in PLA2G15-deficient cells reveal their accumulation in lysosomes and whole cells. (a) Heatmap representation of log2-transformed changes in BMP abundances in lysosomes (IP) and whole cells (WC) of PLA2G15-deficient HEK293T clone1 compared to wildtype control (*n* = 3 WT and *n* = 3 *PLA2G15* KO). Data are the ratio of the mean of each BMP. Significant changes from the graph are represented by *p < 0.05. P values were calculated using two-tailed unpaired tests and are presented in Supplementary Table 1. (b) Fold changes in total BMP abundance. The sum of all abundances of measured BMPs was used to generate these values. Data are mean ± SD. Statistical analysis was performed using two-tailed unpaired *t*-tests. ****p < 0.0001. **(c)** Generation of PLA2G15-deficient clones 2 and 3. (d) Schematic for fluorescently labeled PLA2G15 uptake experiments. Labeled recombinant PLA2G15 proteins were supplemented to *PLA2G15* KO HEK293T cells for 48 hours and imaged to validate delivery. **(e)** Representative images of labeled PLA2G15 wildtype and mutant proteins (Alexa488) and lysosomes (Lysotracker) show successful uptake and trafficking in *PLA2G15* KO HEK293T cells (left). In the merged images, green and magenta represents the protein and Lysotracker channel respectively and white represents the colocalized spots. Scale bar = 5µm. Intensity showing that labeled proteins colocalize with Lysotracker (right). **(f)** Recombinant PLA2G15 does not rescue elevated levels of around a fifth of BMPs resulting from PLA2G15 loss. Fold changes in the levels of BMPs in *PLA2G15* KO HEK293T cells after supplementation with wildtype or mutant PLA2G15 (*n* = 3 WT, *n* = 3 *PLA2G15* KO, *n* = 3 *PLA2G15* KO +PLA2G15 and *n* = 3 *PLA2G15* KO +PLA2G15 S198A). Data are mean ± SD. Statistical analysis was performed using two-tailed unpaired *t*-tests. **p < 0.01, ***p < 0.001 and ****p < 0.0001. **(g)** Targeted analyses of BMP lipids reveal that a deficiency in PLA2G15 increases levels of most BMPs in HeLa cells. Data are mean ± SD (right) (*n* = 3 WT and *n* = 3 *PLA2G15* KO). Statistical analysis was performed using two-tailed unpaired *t*-tests. *p < 0.05, **p < 0.01, ***p < 0.001 and ****p < 0.0001. Individual BMP species and their statistics are presented in Supplementary Table 3. **(h)** Targeted analyses show minimal phospholipid alterations in PLA2G15-deficient mouse tissues (top), HEK293T cells and lysosomes (bottom). Heatmap representation of log2-transformed changes in total lipid class in PLA2G15-deficient mouse tissues (mouse brain, kidney and liver) compared to their control counterparts (*n* = 6 WT and *n* = 6 *Pla2g15*^−/−^) and PLA2G15-deficient HEK293T cells compared to wildtype cells (*n* = 3 *PLA2G15* KO and *n* = 3 WT). Data are the ratio of the mean of each lipid class. Significant changes from the graph are represented by *p < 0.05. P values were calculated using two-tailed unpaired tests. All measured individual lipid species, total lipid classes and their statistics are presented in Supplementary Table 5.**(i and j)** Loss of PLA2G15 has no effect on BMP synthesis and lysosomal biogenesis. (i) Immunoblot analysis of lysosomal markers (LAMP2 and Cathepsin B), the BMP synthase CLN5 and a loading control (Vinculin). (j) BMP synthesis in HEK293T cells using labeled phosphatidylglycerol (D5-PG) at the indicated time points. *n* = 3 WT and *n* = 3 *PLA2G15* KO. Data are mean ± SD.

**Extended Data Fig. 7:**
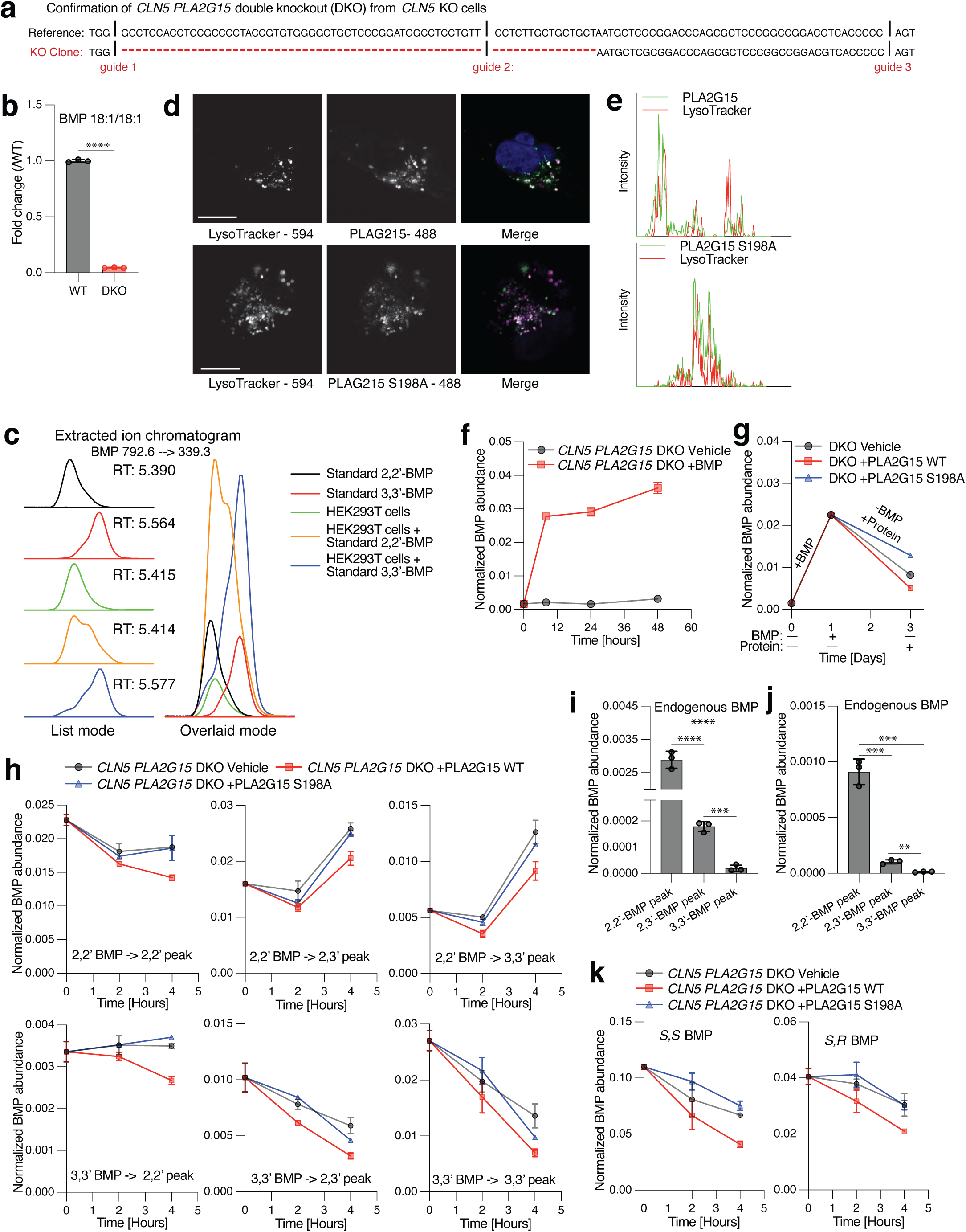
Esterification position in BMP confers resistance to hydrolysis in cells. **(a)** Generation and sequence validation of *PLA2G15* KO on a background of *CLN5* KO HEK293T cells (DKO). **(b)** BMP signal is diminished in *CLN5 PLA2G15* double KO HEK293T cells compared to wildtype. Fold changes of normalized BMP 18:1/18:1 abundance. Data are mean ± SD. Statistical analysis was performed using two-tailed unpaired *t*-tests. ****, p < 0.0001. **(c)** Supplementation of synthesized positional isomers to HEK293T cells demonstrates 2,2’ BMP as the major BMP form in cells. Representative smoothened graphs of the extracted ion chromatograms following addition of 2,2’ and 3,3’ BMP standards to *CLN5 PLA2G15* double KO HEK293T cells. The retention times are shown in the list mode (left) while the alignment of standards in overlaid mode (right) indicate 2,2’ BMP as the predominant endogenous form. **(d)** Representative images of labeled PLA2G15 wildtype and mutant proteins (Alexa488) and lysosomes (Lysotracker) show successful uptake and trafficking in *CLN5 PLA2G15* KO HEK293T cells. In the merged images, green and magenta represents the protein and Lysotracker channel respectively and white represents the colocalized spots. Scale bar = 5µm. **(e)** Curve represents intensity showing that labeled proteins (green) colocalizes with Lysotracker (red) as marker for the lysosome. **(f)** Exogenous BMP feeding to cells was optimized for pulse-chase experiments. Normalized BMP abundance was measured following addition of 3,3’ BMP at indicated time points. **(g)** PLA2G15 WT hydrolyzes BMP in cells while cells supplemented with inactive PLA2G15 S198A exhibit reduced BMP hydrolysis capacity. 100 nM PLA2G15 proteins were supplemented in the chase period after delivery of 10 µM 2,2’ *S,S* BMP for two hours. **(h-j)** Turnover of 3,3’ *S,S* BMP is rapid while 2,2’ BMP may require positional isomerization for efficient degradation in cells. **(h)** The intensity of each BMP peak (2,2’ BMP peak; left, 2,3’ BMP peak; middle, 3,3’ BMP peak; right) was quantified following a 4-hour chase by PLA2G15 proteins or no enzyme control following addition of either 10 µM of 2,2’ *S,S* BMP (top) or 3,3’ *S,S* BMP (bottom). PLA2G15 quickly hydrolyzes 3,3’ and 2,3’ BMP after 3,3’ BMP supplementation while 2,2’ BMP is converted to 2,3’ and 3,3’ BMP. **(i)** Quantitation of endogenous BMP peaks during pulse-chase experiment in cells that were not supplemented with BMP in (h). These levels are minimal compared to cells supplemented with the lipids. **(j)** Quantitation of endogenous BMP peaks in cells that were not supplemented with BMP during pulse-chase experiment (Fig. 4d-f) using equimolar mixture of 2,2’ BMP and 3,3’ BMP. **(k)** BMP stereoisomers are degraded similarly in cells. Same experiment as in (h) using *S,S* BMP (left) and *S,R* BMP (right) to measure total BMP levels during chase.

**Extended Data Fig. 8:**
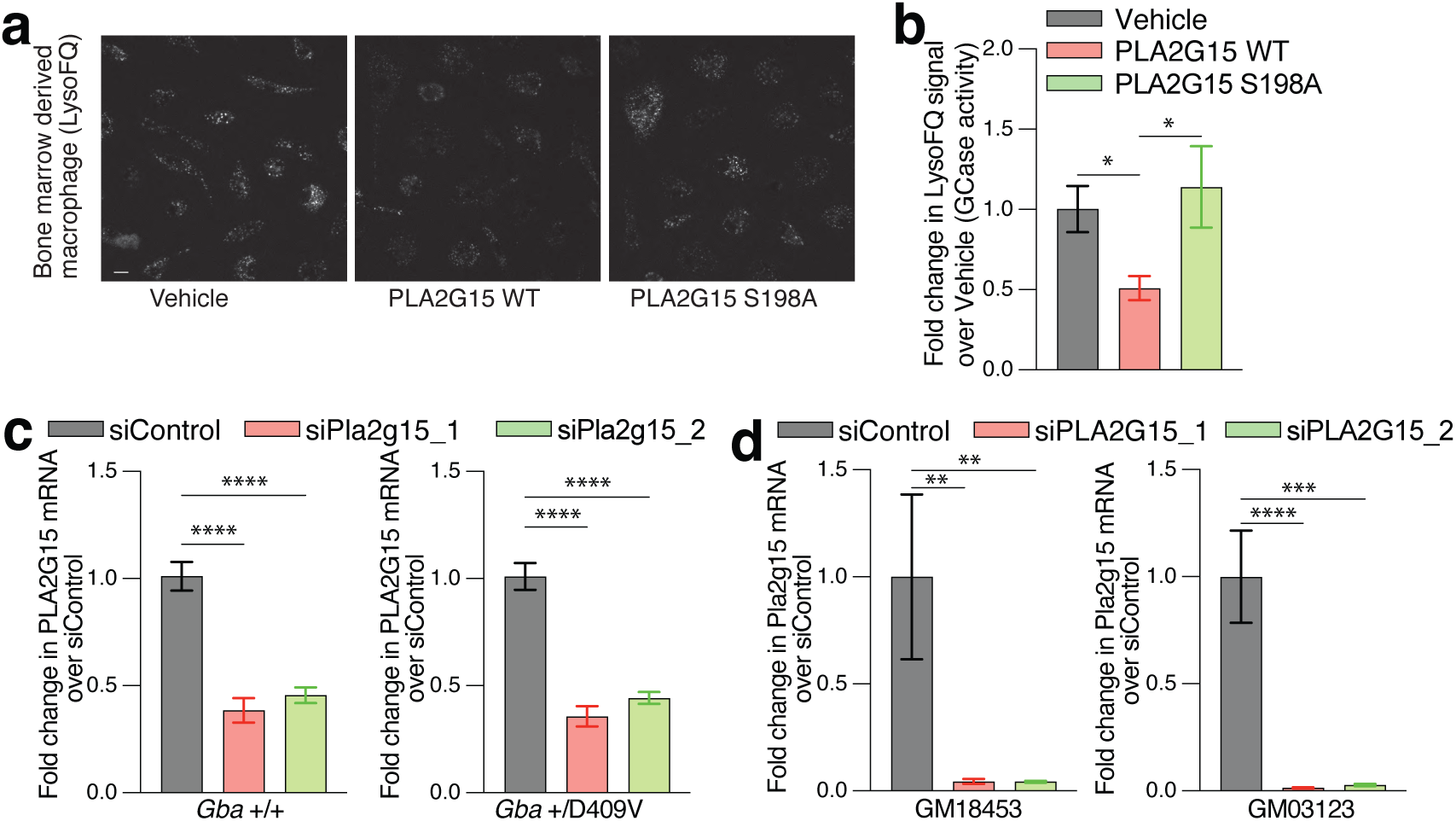
PLA2G15 role in lysosomal lipid metabolism. **(a and b)** PLA2G15 supplementation reduces GCase activity in bone marrow derived macrophages (BMDMs). **(a)** Representative fluorescence microscopy images of BMDMs treated with 300 nM recombinant PLA2G15 WT, inactive PLA2G15 S198A or vehicle buffer (PBS) overnight followed by a 30-minute incubation of 10 µM LysoFQ-GBA to measure GCase activity. **(b)** Quantification of signal intensity of LysoFQ-GBA. Signals were normalized to vehicle treated BMDMs. *, p < 0.05, by one-way ANOVA. **(c)** Pla2g15 knockdown in wildtype and mutant *Gba* mouse-derived BMDMs after two-day treatment with 10 nM control siRNA or two different siRNA that target Pla2g15. qPCR quantification of Pla2g15 mRNA abundance relative to siControl. Actin was used an endogenous control. ****, p < 0.0001, by one-way ANOVA. **(d)** PLA2G15 knockdown in two independent NPC1 patient fibroblast cell lines (Coriell GM03123 and GM18453) after two-day treatment with 10 nM control siRNA or two different siRNA that target PLA2G15. qPCR quantification of PLA2G15 mRNA abundance relative to siControl. GAPDH was used as endogenous control. ****, p < 0.0001, by one-way ANOVA.

**Extended Data Fig. 9:**
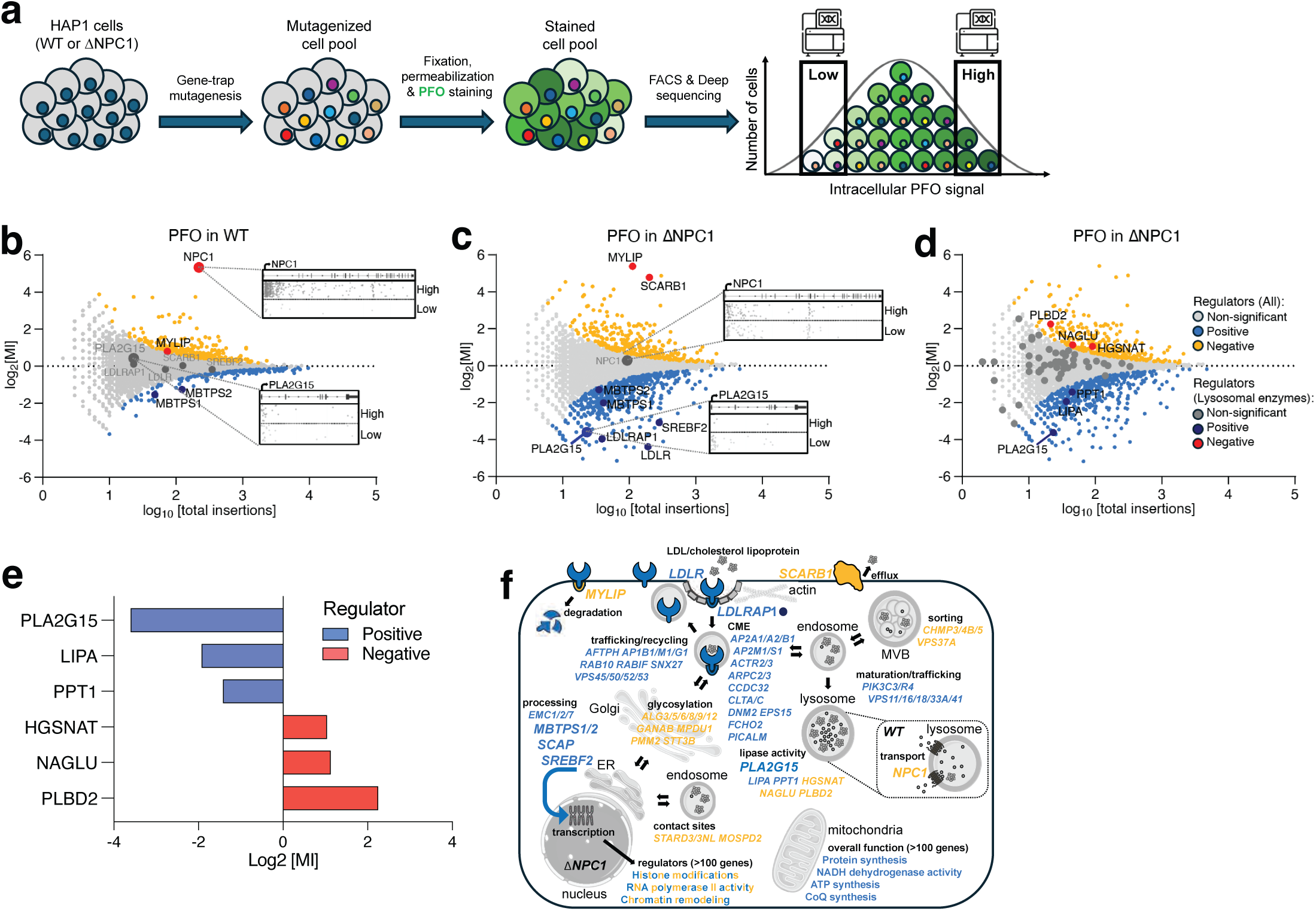
Haploid genetic screens for genes affecting intracellular cholesterol staining. **(a)** Schematic overview of the haploid genetic screens in Hap1 cells. Mutagenized cell libraries were stained for intracellular cholesterol levels using fluorescently labeled PFO. Subsequently, cells are sorted by flow cytometry to obtain cell populations with high and low levels of PFO signal. PFO-HIGH versus PFO-LOW represent the 5% of cells with the highest and lowest fluorescent signal, respectively. **(b and c)** Fishtail plots depicting genetic modifiers of intracellular cholesterol in screens of wild-type (b) and NPC1-deficient (c) Hap1 cells. Insertion sites were mapped in each individual population of cells (i.e. PFO-HIGH versus PFO-LOW) and the log mutational index (MI; see Methods) was plotted against the number of trapped alleles per gene. Statistically significant (p < 0.05) positive (whose loss decreases PFO stain) and negative (whose loss increases PFO stain) regulators are coloured blue and orange, respectively. The screen in wild-type Hap1 cells (b) identified NPC1 as a strong negative regulator of cholesterol staining as expected (in red). In contrast, the screen in NPC1-deficient cells (c) identified genes associated with cholesterol uptake and efflux as strong regulators (i.e. MYLIP, LDLR, LDLRAP1, SREBF2, SCAP, MBTPS1, MBTPS2, SCARB1). In addition, PLA2G15 was identified as a genetic modifier of cholesterol staining (in blue). Note that PLA2G15 was not identified to affect the phenotype in wildtype Hap1 cells. Individual gene-trap insertions (dark grey dots) and their distribution across the gene bodies in PFO-HIGH and PFO-LOW channels of both screens are shown for NPC1 and PLA2G15. **(d)** Same as c but lysosomal genes are highlighted. Out of 55 lysosomal enzyme-encoding genes, 6 appear to regulate cholesterol staining in NPC1-deficient cells. The non-significant genes are labeled dark grey (n=49), while the positive (n=3) and negative (n=3) regulators are labeled in dark blue and red, respectively. **(e)** Of the 6 identified lysosomal enzyme genes, PLA2G15 appears to be the strongest positive regulator of the cholesterol phenotype in NPC1-deficient cells. The other genes include LIPA (Lipase A, lysosomal acid type), PPT1 (Palmitoyl-protein thioesterase 1), NAGLU (N-acetyl-alpha-glucosaminidase), HGSNAT (Heparan-alpha-glucosaminide N-acetyltransferase), and PLBD2 (Phospholipase B Domain Containing 2). The list of 55 enzyme genes can be found in Supplementary Table 6 together with all other screen data. **(f)** Visualization of top hits with known functions in cellular pathways according to GeneCards. Positive and negative regulators are coloured blue and orange, consistent with the fishtail plots in b and c. Positive regulators (in blue) include genes affecting clathrin-mediated endocytosis (CME), receptor trafficking/recycling, endosome maturation/trafficking, and mitochondrial function. Negative regulators (in orange) include genes affecting glycosylation, organelle contact sites, and cargo sorting. Key regulators are indicated with larger font size.

**Extended Data Fig. 10:**
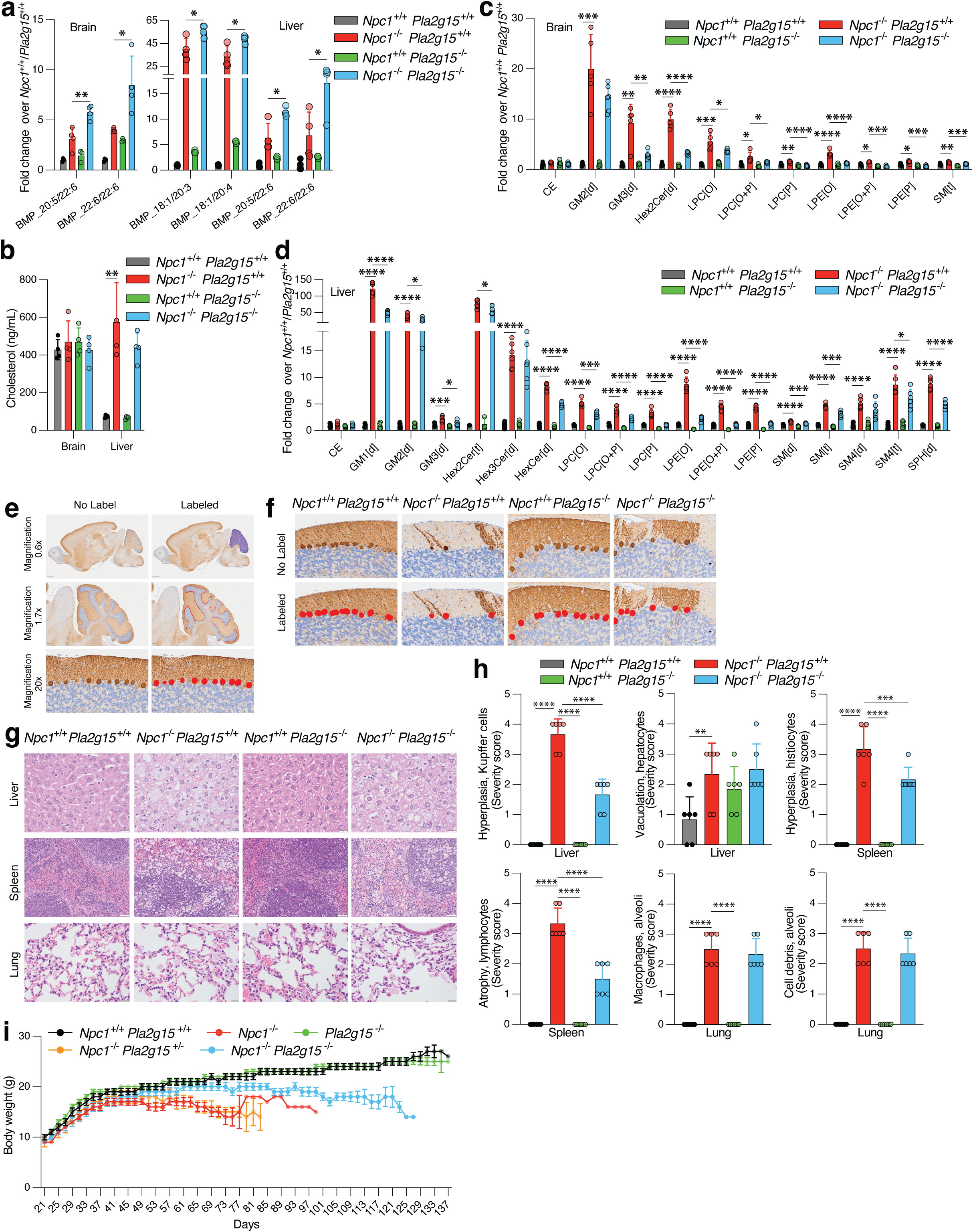
Genetic targeting of *Pla2g15* ameliorates defects in NPC1-deficient mouse. **(a)** PLA2G15 depletion increases levels of only few BMPs in NPC1-deficient mice. Fold changes in the levels of selected BMPs in all genotypes over *Npc1^+/+^ Pla2g15^+/+^* brain (left) and liver (right). Data are mean ± SD (*n* = 4, for all genotypes except *n* = 3 for *Npc1^+/+^ Pla2g15^−/−^*). Statistical analysis was performed using two-tailed unpaired *t*-tests.*, p < 0.05. All measured BMP species and their statistics are presented in Supplementary Table 7. **(b)** Cholesterol levels. Cholesterol was measured in left hemibrain and left liver lobe (*n* = 4 per group). Data are mean ± SD. Statistical analysis was performed using one-way ANOVA followed by Bonferroni’s multiple comparisons. *** p < 0.001. **(c and d)** Genetic depletion of PLA2G15 reverses secondary lipid storage in NPC1-deficient brain (c) and liver (d) mouse tissues. Fold changes in the levels of selected total lipid classes in all genotypes compared to *Npc1^+/+^ Pla2g15^+/+^* brain (c) and liver (d). Data are mean ± SD (*n* = 5, for all brain genotypes except *n* = 6 for brain *Npc1^−/−^ Pla2g15^−/−^* and *n* = 6, for all liver genotypes). Statistical analysis was performed using two-tailed unpaired *t*-tests.*, p < 0.05. All measured individual lipid species, total lipid classes and their statistics are presented in Supplementary Table 8. **(e and f)** Purkinje cell count by histomorphometry on Calbindin-immunolabelled CNS sections. (e) Proof-of-concept of the image analysis indicating the annotations in the cerebellum (ROI: Region of Interest) and the detection of Purkinje cell soma at different magnifications. **(f)** Representative images of the cerebellum at the level of lobe II/III for comparison among study groups. No label: Scanned whole slide images (WSI) prior to image analysis. Labeled: WSI indicating the annotations of the ROI and the detections of Purkinje cell’s soma. **(g and h)** Genetic ablation of PLA2G15 decreases histopathology lesions in NPC1-deficient mice. The microscopy images are in g and their mean severity scores of each histopathology finding are shown in h. Data are mean ± SD (n = 6) and **, p < 0.01, ****, p < 0.0001, by one-way ANOVA comparing NPC1-deficient tissues to other genotypes. Scale bar is 20 µm. **(i)** Body weight assessment for all genotypes. Animal body weight was measured every other day starting at 3 weeks old. Data are mean ± SEM.

### Supplementary Tables

**Supplementary Table 1**: Targeted BMP analyses of lysosomes and whole cells from PLA2G15-deficient and wildtype HEK293T cells.

**Supplementary Table 2**: Targeted BMP analyses from wildtype and PLA2G15-deficient HEK293T cells.

**Supplementary Table 3**: Targeted BMP analyses from PLA2G15-deficient and wildtype HeLa cells.

**Supplementary Table 4**: Targeted BMP analyses from PLA2G15-deficient mouse tissues and their control counterparts.

**Supplementary Table 5**: Targeted lipidomic analyses from PLA2G15-deficient mouse tissues, cells and lysosomes, and their control counterparts.

**Supplementary Table 6**: Haploid genetic screening data for PFO staining in wildtype and NPC1-deficient cells.

**Supplementary Table 7**: Targeted BMP and Hemi-BMP analyses from NPC1-deficient or/and PLA2G15-deficient mouse tissues and their control counterparts.

**Supplementary Table 8**: Untargeted lipidomic analyses from NPC1-deficient or/and PLA2G15-deficient mouse tissues and their control counterparts.

**Supplementary Table 9**: CNS Histopathological Evaluations of NPC1-deficient or/and PLA2G15-deficient mouse tissues and their control counterparts.

**Supplementary Table 10**: Multiple reaction monitoring (MRM) transitions table for targeted lipid and metabolite analyses.

